# Type-I interferon signatures in SARS-CoV-2 infected Huh7 cells

**DOI:** 10.1101/2021.02.04.429738

**Authors:** Xi Chen, Elisa Saccon, K. Sofia Appelberg, Flora Mikaeloff, Jimmy Esneider Rodriguez, Beatriz Sá Vinhas, Teresa Frisan, Ákos Végvári, Ali Mirazimi, Ujjwal Neogi, Soham Gupta

## Abstract

Severe acute respiratory syndrome coronavirus 2 (SARS-CoV-2) that causes Coronavirus disease 2019 (COVID-19) has caused a global health emergency. A key feature of COVID-19 is dysregulated interferon-response. Type-I interferon (IFN-I) is one of the earliest antiviral innate immune responses following viral infection and plays a significant role in the pathogenesis of SARS-CoV-2. In this study, using a proteomics-based approach, we identified that SARS-CoV-2 infection induces delayed and dysregulated IFN-I signaling in Huh7 cells. We demonstrate that SARS-CoV-2 is able to inhibit RIG-I mediated IFN-β production. Our results also confirm the recent findings that IFN-I pretreatment is able to reduce susceptibility of Huh7 cells to SARS-CoV-2, but not post-treatment. Moreover, senescent Huh7 cells, in spite of showing accentuated IFN-I response were more susceptible to SARS-CoV-2 infection, and the virus effectively inhibited IFIT1 in these cells. Finally, proteomic comparison between SARS-CoV-2, SARS-CoV and MERS-CoV revealed a distinct differential regulatory signature of interferon-related proteins emphasizing that therapeutic strategies based on observations in SARS-CoV and MERS-CoV should be used with caution. Our findings provide a better understanding of SARS-CoV-2 regulation of cellular interferon response and a perspective on its use as a treatment. Investigation of different interferon stimulated genes and their role in inhibition of SARS-CoV-2 pathogenesis may direct novel antiviral strategies.

## Introduction

The novel severe acute respiratory syndrome coronavirus 2 (SARS-CoV-2) that emerged in the end of 2019, caused a major ongoing pandemic with more than a million deaths worldwide by the end of 2020 ^1^. SARS-CoV-2 belongs to the genus betacoronavirus, which also includes SARS-CoV and MERS-CoV, two viruses that caused outbreaks in 2002 and 2012, respectively ^2^. These viruses have the capability of infecting both upper and lower respiratory tract with potential to cause severe and fatal respiratory syndrome in humans ^3^. While SARS-CoV-2 presented a lower case-fatality than SARS-CoV and MERS-CoV, they shared similar clinical features ^2,4^. The severe form of the disease is often associated with a dysregulated type-I interferon (IFN-I) response leading to the pathogenesis ^5^, which is attributed to the immunomodulatory proteins encoded by the coronaviruses.

Type-I interferon response that majorly constitutes IFNα and IFNβ is produced by almost every cell and is one of the first lines of defense against viruses ^6^. The early activation of IFN responses against coronaviruses is initiated by recognition of viral products by the host pattern recognition receptors like Toll-like receptors (TLRs) and RIG-I like receptors (RLRs). RLRs can recognize the viral RNA that promotes their oligomerization and subsequent activation of a signaling cascade leading to production of IFNα and IFNβ ^7^. Through autocrine and paracrine signaling the secreted IFN can bind to IFN-α/β receptors (IFNARs) that activates the Janus kinase 1 (JAK1) and Tyrosine kinase 2 (Tyk2) leading to phosphorylation of signal transducer and activator of transcription proteins, STAT1 and STAT2. Activated STAT1 and STAT2 form the interferon-stimulated gene factor 3 (ISGF3) complex in association with IRF9, translocate to the nucleus with the help of nuclear transporter proteins, bind to IFN-stimulated response elements and trigger transcription of several interferon stimulated genes (ISGs) with antiviral properties ^8^. Coronaviruses also have evolved mechanisms to evade the host’s antiviral immune response. Several proteins in SARS-CoV (nsp1, PLpro, nsp7, nsp15, ORF3b, M, ORF6 and N) ^9^, in MERS-CoV (M, ORF4a, ORF4b, PLpro and ORF5) ^3,9^ and in SARS-CoV-2 (ORF6, PLpro, nsp6, nsp13, nsp1, ORF3a, ORF7a/b and M) ^10,11^ have been shown to be strong IFN-antagonist.

Most of the current treatment options for SARS-CoV-2 have been guided by knowledge on SARS-CoV and MERS-CoV infection. Based on which therapeutic interventions with type-I IFN treatment and remdesivir have been employed for SARS-CoV-2 ^12–14^. However, the dynamics of the IFN response in mouse models of SARS-CoV and MERS-CoV was observed to vary ^15,16^ as well as the sensitivity to IFN-treatment *in vitro* ^17^. Moreover, the transcriptome analysis comparing *in vitro* host cell response to SARS-CoV-2, SARS-CoV and MERS-CoV have shown distinct virus specific patters ^18^. Thus, a deeper understanding of the SARS-CoV-2 mediated regulation of IFN response is necessary to develop rationale and novel therapeutic approaches for SARS-CoV-2

In this study, we characterized the SARS-CoV-2 mediated dysregulation of IFN-signaling in Huh7 infected cells using quantitative proteomics. We show a delayed activation of IFN-signaling with the ability of the virus to evade RIG-I mediated IFN-signaling during early infection. In line with recent studies susceptibility of Huh7 cells to SARS-CoV-2 decreased upon IFN-pretreatment, but not post-treatment. We also determined the IFN-signaling response pattern of SARS-CoV and MERS-CoV infection in Huh7 cells using proteomics and show a distinction compared to SARS-CoV-2. Together, the results provide a perspective of immune regulation by coronaviruses.

## Results

### Quantitative proteomics and transcriptomics of SARS-CoV-2 infected Huh7 cells identifies dysregulation in type-1 interferon signaling pathway

Interferons (IFNs) play a critical role in exerting an early antiviral response to inhibit viral replication and spread. To understand how the IFN responses are modulated following SARS-CoV-2 infection, we re-used the proteomics and transcriptomics data set from our earlier study ^19^. We first analyzed the quantitative proteomics data on Huh7 cells that were either mock infected or infected with SARS-CoV-2 at multiplicity of infection (MOI) of 1, over a period of 24 and 48 hours post infection (hpi). Genes associated with the interferon response, including the interferon alpha/beta signaling (Pathway: R-HSA-909733), interferon gamma signaling (Pathway: R-HSA-877300) and antiviral mechanism by IFN-stimulated genes (ISGs, Pathway: R-HSA-1169410) were extracted from the data. For mock infected we considered the data for two replicates as one of the replicated was a major outlier as shown in the PCA plot (Supplementary Figure 1). No major changes were observed in the interferon signaling genes at 24hpi and significant modulation was only observed at 48hpi after infection as represented in the heatmap (Figure 1A). Of the 94 proteins studied, a number of proteins showed significant reduction in abundance (n=20), while a major cluster of proteins showed an increase (n=26) (LIMMA, FDR < 0.05). The log2 foldchange of the significantly regulated genes are represented as volcano plot in Figure 1B. The protein-protein interaction network of the significantly changed genes showed two definite clusters (cluster-1 and cluster-2). Cluster-1 involved proteins associated with the RIG-I (DDX58) and type-1 interferon signaling cascade, while cluster-2 mostly involved proteins associated with transporters belonging to the components of nucleoporin complex and karyopherin family (Figure 1C).

**Figure 1:**
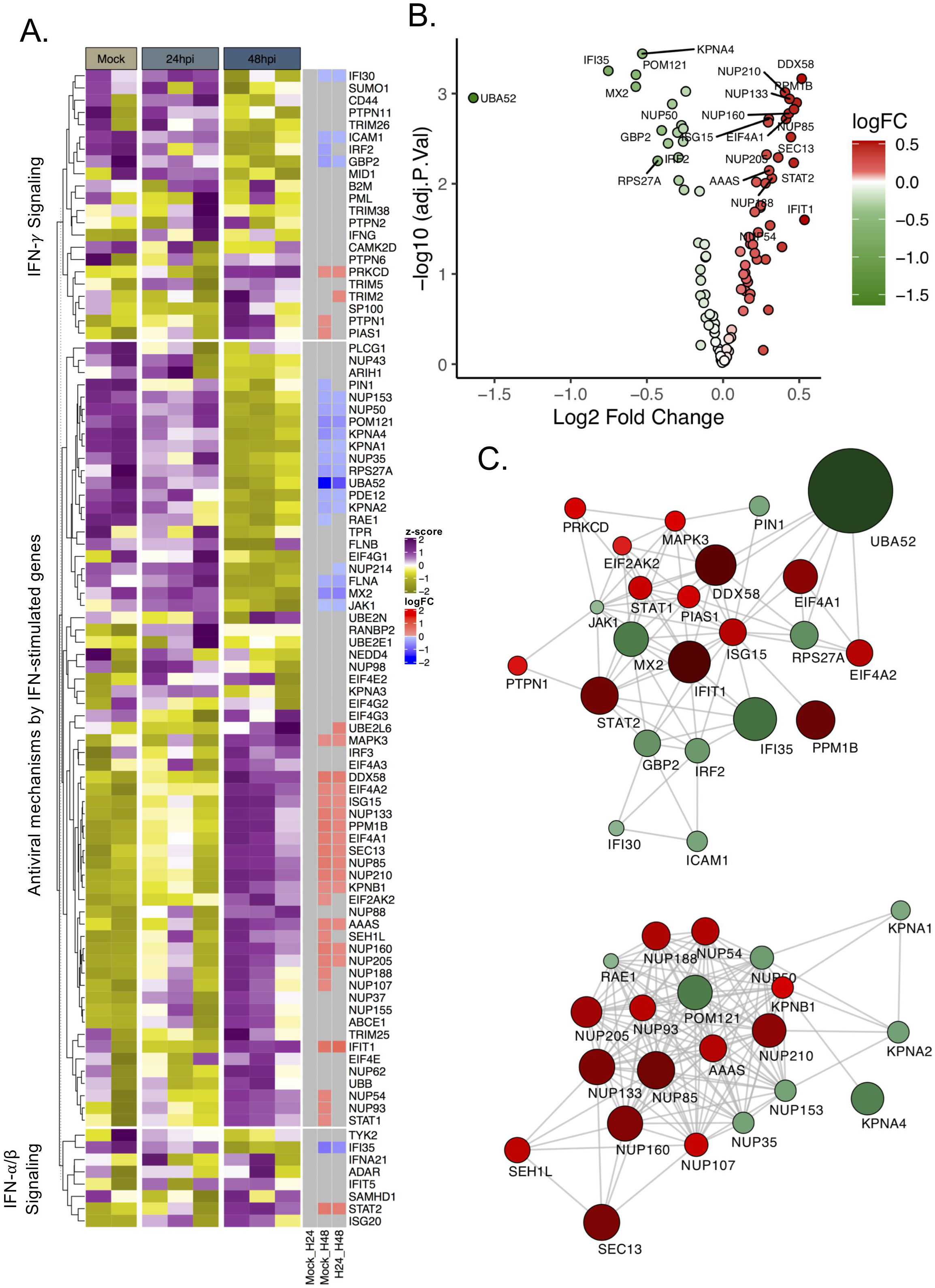
SARS-CoV-2 induced a delayed and dysregulated IFN signaling response identified in proteomics data. A) Heatmap of IFN-stimulated proteins before infection and at 24 and 48hpi. Data were quantile normalized and Z-score transformed. Lower values are represented in yellow and higher values in purple. Significant differential expressed proteins between time points are indicated in blue if downregulated and in red if up regulated. B) Volcano plots of proteins with differential abundance between Mock and Huh7 cell 48 h after SARS-CoV-2 infection. Upregulated proteins are represented in red while proteins downregulated are represented in green. FDR < 0.05. C) Cytoscape network of differentially abundant IFN-stimulated proteins. Proteins are represented as circles. Gradient color was applied on proteins depending on fold change (low = green to high = red). Size of the circle is proportional to the fold change.

We also looked into the IFN-signaling genes in the transcriptomics dataset and observed no major changes in the differential expression of the transcripts related to this pathway except for EIF4A2, STAT2, TRIM10 (upregulated) and FLNA, JAK1, GBP2, MT2A, TRIM26 (down regulated) 48 h after infection (Figure 2A). Of the genes corresponding to the proteins that were altered in the pathway (Figure 2B) only EIF4A2, STAT2, JAK1, GBP2 and FLNA showed transcript levels correlating with protein expression (Figure 2C). Of note, we had previously observed major changes in the global transcriptome to occur only after 72 h of infection ^19,20^.

**Figure 2:**
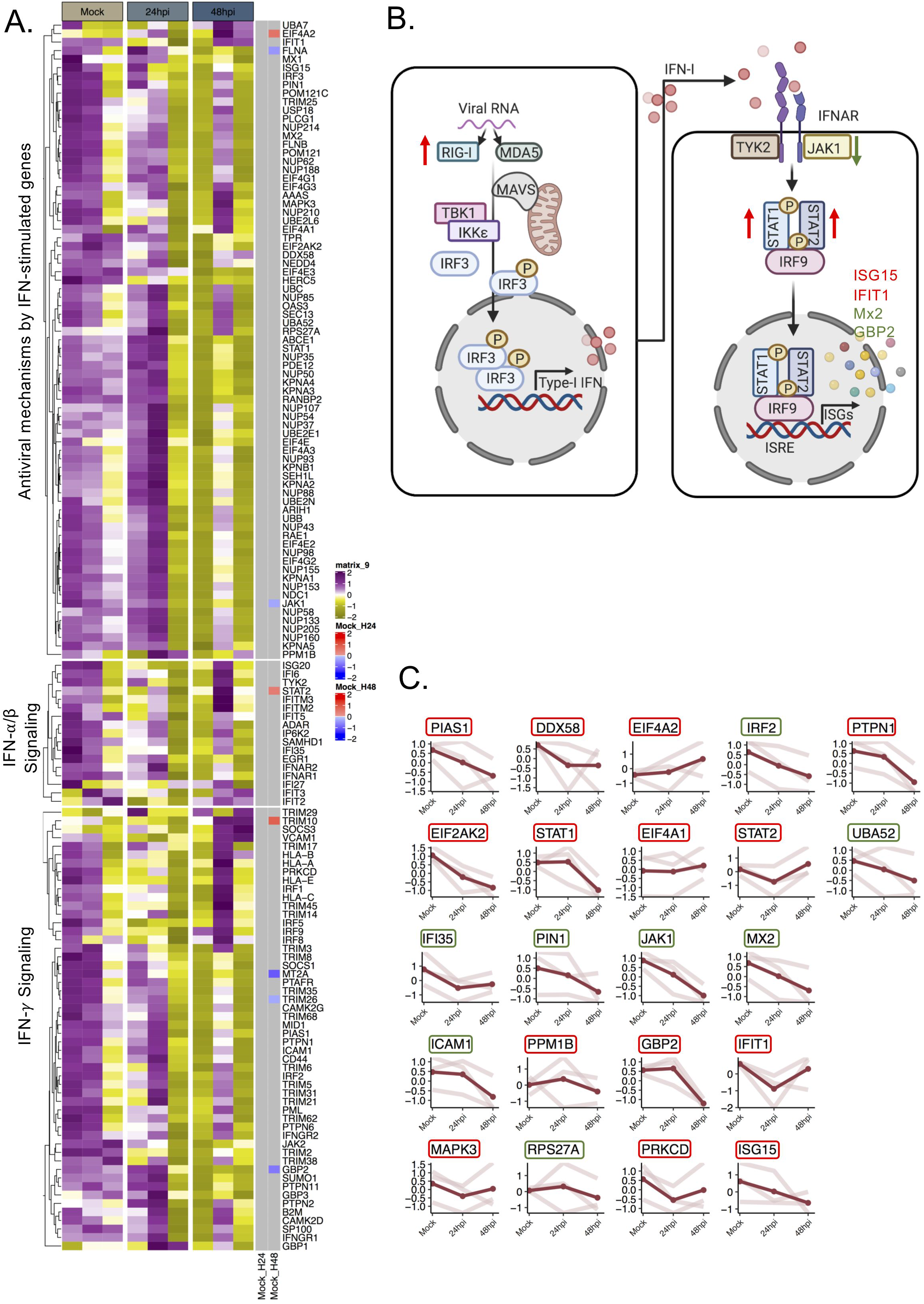
SARS-CoV-2 induced transcriptional changes in the IFN-signaling genes in transcriptomics data. A) Heatmap of IFN-stimulated transcripts before infection and at 24 and 48hpi. Data were log2 normalized and Z-score transformed. Lower values are represented in yellow and higher values in purple. Significant differential expressed genes between time points are indicated in blue if downregulated and in red if up regulated. B) The scheme graph of the type I interferon signaling pathways, in which the regulated genes expression level trend is noted. The significantly changed proteins observed in the proteomics data are denoted by green arrows or letters (downregulated) or red arrows or letters (upregulated) C) Dot plot for each transcript that were detected as significantly altered in proteomics. For each gene, the scaled values in triplicates are represented in mock, 24hpi and 48hpi and linked by light red line, average value is displayed in red. The name of the genes is indicated in colored box based on the proteomics data. The genes corresponding to increased protein levels are in red boxes and to decreased protein levels in green boxes.

### SARS-CoV-2 induces delayed and low-level activation of RIG-I signaling in Huh7 cells

In our global transcriptomics and proteomics data we observed a delayed activation of RIG-I and dysregulation of type-I IFN response associated proteins including ISGs. RIG-I, a key cytosolic receptor that can detect SARS-CoV-2 RNA is responsible for activation of IFN-β through a signaling cascade that can further lead to activation of antiviral ISGs (Figure 3A). We next studied the effect of SARS-CoV-2 in induction of IFN-β. We did not observe any significant changes in the levels of IFN-β specific mRNAs in SARS-CoV-2 infected Huh7 cells both at 24hpi and 48hpi (Figure 3B). Even though not significant, we observed an increase in IFN-β mRNA at 48hpi with an infective dose of MOI 0.1 (Figure 3B). This effect was concomitant with a marginal suppression of RIG-I and MDA-5 protein expression at 24 hpi and an observable increase in at 48 hpi detected in western blots probed with specific antibodies (Figure 3C and 3D). The Western blot data was in line with our proteomics data.

**Figure 3:**
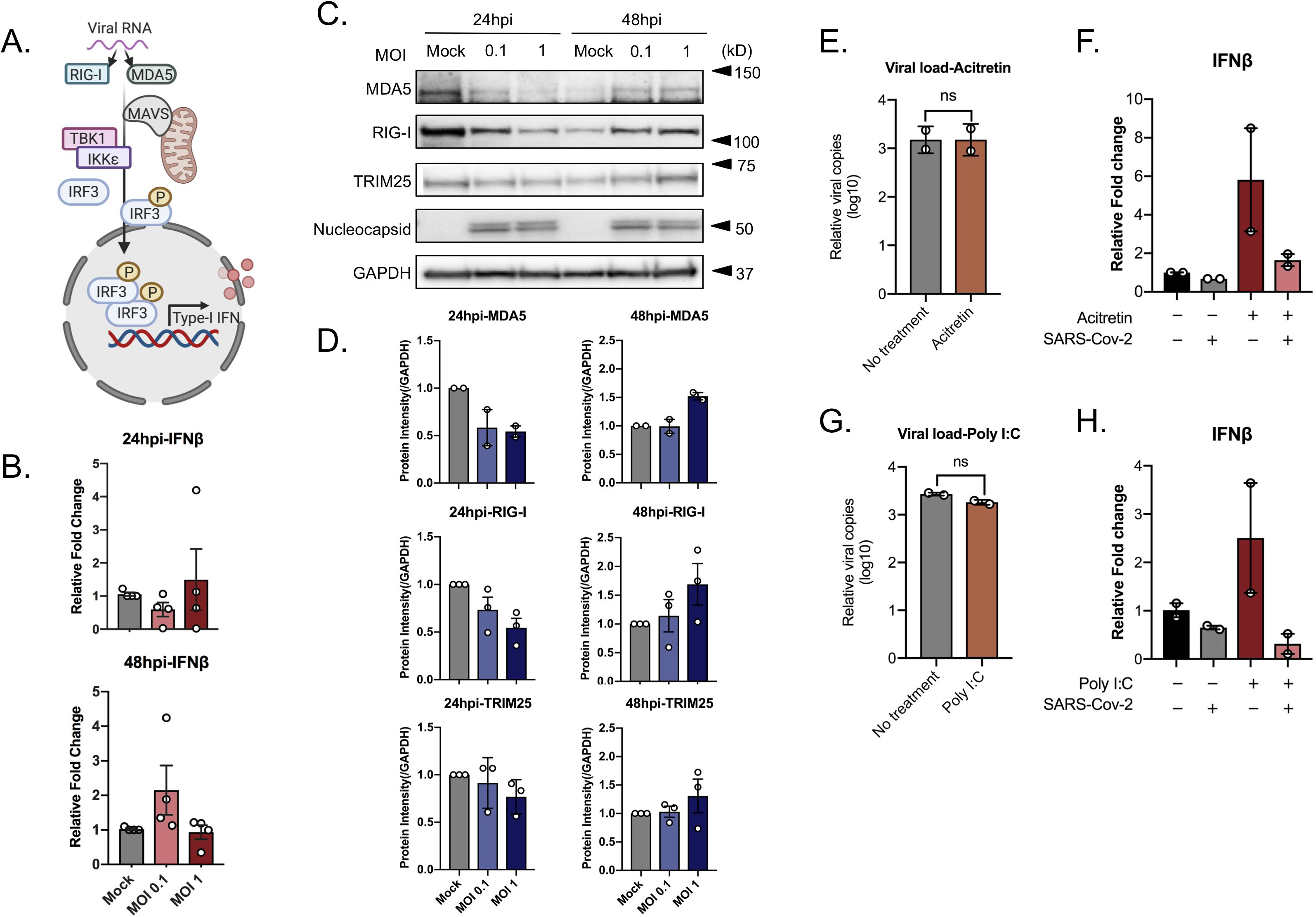
SARS-CoV-2 induces delayed and low-level activation of RIG-I signaling and antagonizes IFN-β activation. To check the RIG-I signaling response on SAR-CoV-2 infection, Huh7 cells were infected with SARS-CoV-2 at MOI of 0.1 and 1. Cells were collected at 24hpi and 48hpi. To assess if SARS-CoV-2 can inhibit the activation of IFN-β, Huh7 cells were infected with SARS-CoV-2 at MOI 0.1 treated with acitretin (25 μM) or poly I:C (5 μg/mL) 16 h before infection. Cells and cell supernatant were harvested at 24hpi. Virus production in the cell culture supernatant was determined by quantitative RT-PCR targeting the E-gene of SARS-CoV-2. An unpaired t-test was used to determine p-values (ns, *p > 0.05*) A) Schematic representation of RIG-I/MDA-5 signaling pathways. B) IFN-β transcripts level in SARS-CoV-2 infected (MOI 0.1 and MOI 1) or mock infected cells were quantified by qRT-PCR, normalized to GAPDH as a reference gene. The results are shown as fold change relative to mock treated cells. The mean ± SEM of four independent experiments is shown. C) Western blots of the cell lysates were probed with the indicated antibodies. One representative experiment out of 3 is shown. D) The intensity of specific bands was quantified by ImageJ and fold change was calculated relative to the uninfected cells(mock), normalized to GAPDH. The mean ± SEM of at least two experiments is shown. E) Production of the virus following acitretin treatment. The mean ± SD of two independent experiments is shown. F) IFN-β transcripts level following acitretin treatment. The mean ± SEM of two independent experiments each performed in duplicate is shown. G) Production of the virus following poly I:C treatment. The mean ± SD of two independent experiments is shown. H) IFN-β transcripts level following poly I:C treatment. The mean ± SEM of two independent experiments is shown.

### SARS-CoV-2 can inhibit IFN-β activation

SARS-CoV-2 infection was shown to induce high level of IFN-β in Calu-3 cells and Caco2 cells at 24hpi ^21,22^. However, we did not observe any IFN-β induction or RIG-I activation at 24hpi, suggesting that SARS-CoV-2 is able to inhibit IFN-β activation in Huh7 cells. To determine this Huh7 cells were either mock infected or infected with SARS-CoV-2 at MOI 0.1, followed by either treatment with RIG-I agonist acitretin or transfected with poly I:C for 24h to induce transcription of IFN-β. Treatment with acitretin or poly I:C post-infection did not inhibit production of the virus as measured by qPCR targeting the E-gene in the cell culture supernatant (Figure 3E and 3G). SARS-CoV-2 was able to efficiently inhibit the IFN-β production in the RIG-I activated cells (Figure 3F and 3H).

### SARS-CoV-2 regulates host-protein ISGylation

IFN-β produced by a cell binds to IFN-α/β membrane receptors (IFNAR), activating the JAK-STAT signaling cascade, which leads to expression of several ISGs with antiviral properties (Figure 4A). Similar to our transcriptomics data, qPCR analysis to detect IFIT1, RIG-I (DDX58) and MX2 in SARS-CoV-2 infected Huh7 cells did not show any significant changes in RNA expression of these genes compared to uninfected cells (Figure 4B). Though, in our proteomics data we observed several ISGs to be stimulated among which ISG15 showed an increase in protein level 48hpi. ISG15 can conjugate itself to host proteins to regulate diverse cellular functions as well as viral proteins to alter their mechanisms (Figure 4A) ^23^. In the unconjugated form ISG15 can behave as a cytokine with ability to inhibit viral replication ^24^. We examined the mRNA levels of ISG15 in SARS-CoV-2 infected Huh7 cells after 24h and 48h. We did not observe any significant change in ISG15 at transcript level (Figure 4C). However, at protein level it was interesting to note that there was an observable decrease in the conjugated ISG15 at 24hpi and a marked increase in host-cell ISGylation at 48hpi (Figure 4D and 4E) in a dose dependent manner, suggesting the virus can modulate cellular ISGylation to alter the cellular environment. It was not surprising to observe a decreased ISGylation during early infection as SARS-CoV-2 encodes papain-like protease (PLpro) that is a potent de-ISGylase ^11^.

**Figure 4:**
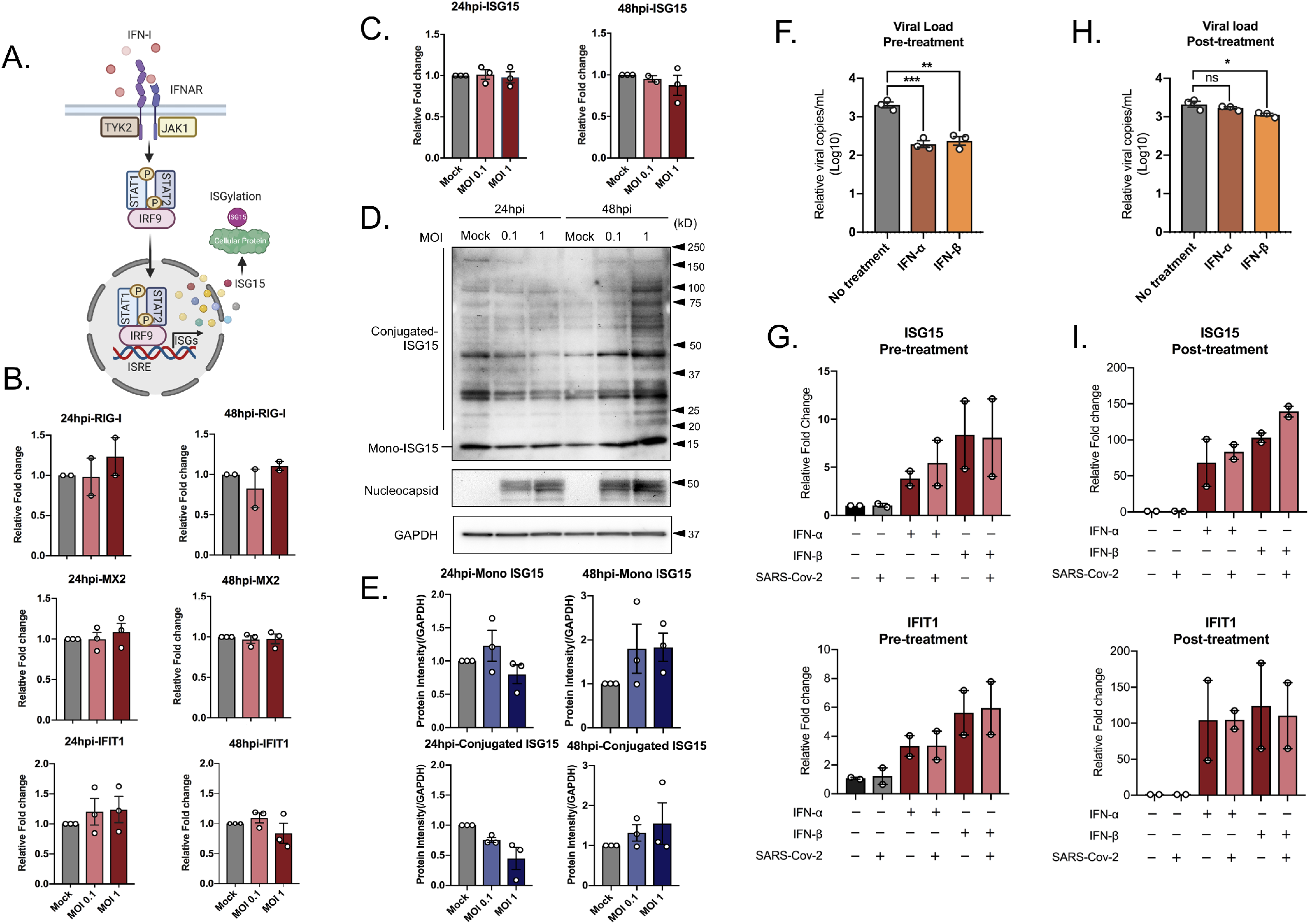
SARS-CoV-2 regulates host-protein ISGylation and is sensitive to IFN pretreatment. To understand the regulation of type I interferon induced signaling pathways, Huh7 cells were infected with SARS-CoV-2 at MOI of 0.1 and 1. Cells were collected at 24hpi and 48hpi. A) Schematic representation of the activation of JAK/STAT pathways and interferon stimulated genes. B) The transcripts expression level of some representative interferon stimulated genes (ISGs): RIG-I, MX2, and IFIT1. The results are shown as fold change relative to mock treated cells, normalized to GAPDH. The mean ± SEM of at least two independent experiments is shown. C) The ISG15 transcript levels. The results are shown as fold change relative to mock treated cells, normalized to GAPDH. The mean ± SEM of three independent experiments is shown. D) ISG15 protein levels in SARS-CoV-2 infected at MOI of 0.1 and 1, or mock infected. The representative western blots with the indicated antibodies are shown. E) The intensity of specific bands was quantified by ImageJ and fold change was calculated relative to the uninfected cells (mock). The mean ± SEM of three experiments is shown. To determine the effect of Type I interferon on SARS-Cov-2 infection, Huh7 cells were treated with 5000 u/mL IFN-α, 100 u/mL IFN-β 16h prior or 24 h after infection. The cells were infected with SARS-CoV-2 at a MOI of 0.1, the mean ± SEM is shown. Unpaired t test was used to determine p-values (* p ≤ 0.05, ** p ≤ 0.01, *** p ≤ 0.001, **** p ≤ 0.0001) F) The virus production in the cell culture supernatant in type I interferon pre-sensitized cells. The mean ± SD of three independent experiments is shown. G) ISG15 and IFIT1 Transcripts level in type I interferon pre-sensitized cells. The mean ± SD of two independent experiments each performed in triplicate is shown. H) The virus production in the cell culture supernatant in post-infection type I interferon treated cells. The mean ± SD of three independent experiments each performed in duplicate is shown. I) ISG15 and IFIT1 transcripts levels were evaluated in response to type I interferon treatment post infection. The mean ± SD of two independent experiments each performed in triplicate is shown.

### SARS-CoV-2 is inhibited by IFN pretreatment

SARS-CoV-2 was observed to change the levels of different ISGs in Huh7 cells (Figure 1A, 1B and 1C). ISGs can also be stimulated in experimental models by external treatment with IFNs. In order to evaluate the susceptibility of SARS-CoV-2 to type-I IFN (IFN-I), we either pre-sensitized cells (16h) with IFN-α (5000u/mL) and IFN-β (100u/mL) or treated the cells with the same concentrations of IFNs starting 1hpi and continued for 24h. Huh7 cells were infected with SARS-CoV-2 at MOI 0.1 and at 24hpi the supernatant was collected to determine the virus production in presence or absence of different IFN-I treatment conditions. As shown in Figure 4F, IFN-pre-sensitization lead to a significant reduction in SARS-CoV-2 production in the supernatant as compared to levels in supernatant from untreated cells at 24hpi.. However, IFN-I treatment after infection did not suppress virus production (Figure 4H). This observation suggests firstly that the presence of high level of IFN-response can suppress the incoming virus and secondly that the virus has also developed measures to counteract these responses when it has already established an infection. Then, we further looked into the effect of IFN-I treatment and infection in transcriptional activation of few of the ISGs that were modulated by SARS-CoV-2 infection. For this we selected MX2, IFIT1 and ISG15. While SARS-CoV-2 suppressed MX2 mRNA in untreated cells, MX2 did not show any activation following IFN-treatment (data not shown). Both ISG15 and IFIT1 were significantly induced following IFN-I treatment, however SARS-CoV-2 did not cause any significant alterations to the mRNA levels (Figure 4G and Figure 4I respectively).

### Senescent Huh7 cells stimulate IFN-I response but promotes virus infectivity

Elderly people has been suggested to be more susceptible to SARS-CoV-2 infection ^25^ and cellular senescence is postulated as factor for increased infection. Cellular senescence has been observed to play different role in either promoting infection for some viruses or inhibiting infection for others. To this end we aimed to examine the susceptibility of senescent Huh7 cells to SARS-CoV-2 and associated IFN-I response. To induce cellular senescence Huh7 cells were treated with 0.5 μM of etoposide for 6 days followed by 2 days without any treatment and then infected with SARS-CoV-2 for 1h and cells and supernatants were harvested 24hpi. Etoposide treatment resulted in massive cell death and surviving cells were large in size. Cellular senescence was determined by detecting p21 mRNA levels (Figure 5B, top-panel leftmost). SARS-CoV-2 infectivity was determined by measuring the viral E-gene in the supernatant. Senescent Huh7 cells showed a significant increase in virus production in senescent Huh7 cells compared to the etoposide untreated control cells (Figure 5A). We next investigated the IFN-response in senescence-induced and non-induced cells by detecting mRNA transcripts of IFN-β and ISG’s such as ISG15, IFIT1, MX2 and RIG-I. Cellular senescence induced an increase in the IFN-response with significant increase in the levels of IFN-β and other ISG’s tested (Figure 5B). Among the genes tested SARS-CoV-2 failed to significantly alter the levels of any except for IFIT1, where a significant decrease in the mRNA levels were noted upon infection (Figure 5B, top-panel second from left). To determine if the enhanced infectivity of senescent cells is specific to Huh7, we tried to replicate the same experiment in Caco2 cells. However, Caco2 cells were more resistant to 0.5 μM etoposide treatment and did not show observable induction of senescence as observed by the qPCR of p21 gene (Supplementary Figure S2B, top-panel leftmost). Most interestingly, in contrast to Huh7, even a very low-level induction of p21 was sufficient to significantly reduce SARS-CoV-2 susceptibility (Supplementary Figure S2A) and among the ISG’s IFIT1 showed an observable increase upon infection (Figure S2B, top-panel second from left). The results suggest that there is a cell-type specific regulation of SARS-CoV-2 and importance of IFIT1 as an anti-SARS-CoV-2 ISGs.

**Figure 5:**
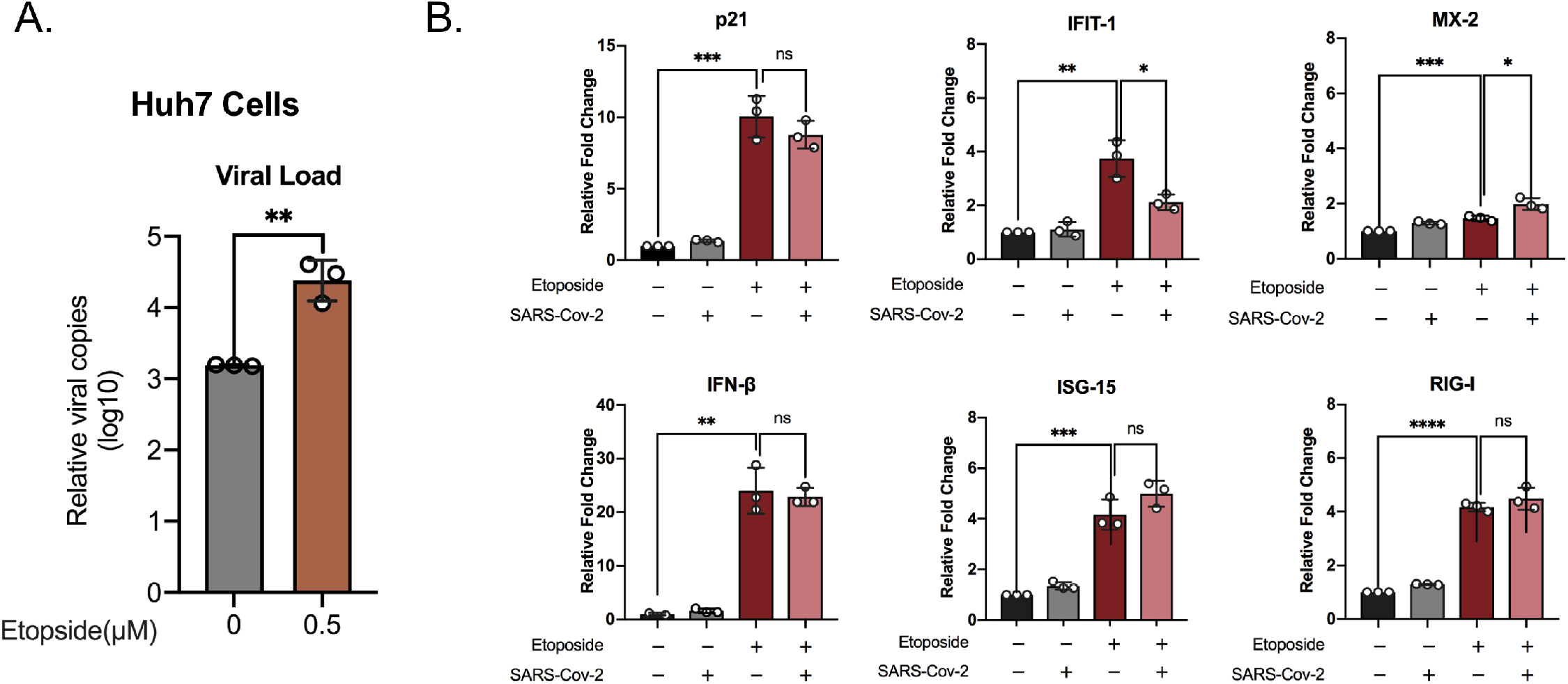
Senescent Huh7 cells shows increased susceptibility to SARS-CoV-2 infection. To determine the susceptibility of senescent Huh7 cells to SARS-Cov-2 and associated IFN-I response, Huh7 cells were treated with 0.5 μM of etoposide for 6 days followed by 2 days regular DMEM with 10%FBS. The cells were either mock infected or infected with SARS-CoV-2 at MOI of 0.1. After 24h the cell-culture supernatant and cells were harvested to determine the virus production and the transcript levels of the indicated genes respectively. The experiments were performed in technical triplicate and the mean ± SD values is shown. An unpaired t-test was used to determine p-values (* <0.05, **< 0.01, ****< 0.001) A) The virus production in senescent Huh7 cells. B) The levels of specific mRNAs were quantified by qRT-PCR. The results are shown as fold change relative to non-treated cells. The mean± SD of technical triplicates are shown.

### Global proteomic response to SARS-CoV-2 relative to SARS-CoV and MERS-CoV in Huh7 cells

To explore the differences in pathogenicity of SARS-CoV-2 in comparison with its predecessor human pathogenic coronaviruses; SARS-CoV and MERS-CoV, we measured the global proteomic changes by infection in the same cell line and at the same infective dose. MERS-CoV-2 was observed to be highly cytopathic and by 48hpi all the cells were dead restricting our analysis to 24hpi, while SARS-CoV showed a slower cytopathogenicity and infected cells were collected both at 24hpi and 48hpi. Quantitative proteomics was performed utilizing a TMT-labeling strategy of mock infected and virus-infected cells in triplicate as previously described by us ^19^. The PCA plots are shown in supplementary Figure S3 and level of infection by the virus in the cells was determined by detecting the increase in viral proteins abundance as shown in supplementary Figure S4. Overall, MERS-CoV infection showed significant differences in 1344 proteins compared to the mock infected (LIMMA, FDR < 0.05), while SARS-CoV showed a significant difference in 165 protein at 24hpi and 310 proteins by 48hpi (LIMMA, FDR < 0.05). We next examined the pathways that were enriched in common proteins with differential abundance in SARS-CoV, MERS-CoV and SARS-CoV-2 infected cells compared to mock using ClusterProfiler. We observed that several pathways in relation to infectious diseases, rRNA processing and mRNA translation were significantly regulated by all the three viruses (Supplementary Figure S5). For the current paper we focused our analysis to regulation of IFN-response. In the IFN-signaling pathways, we looked at proteins that were differentially regulated by any of the three viruses and they are represented as a heatmap in Figure 6A. SARS-CoV showed very little change in IFN-related proteins (n=5) and MERS-CoV showed changes in the levels of 48 proteins. It was interesting to observe that there was no overlap between significantly altered IFN-related proteins in SARS-CoV and MERS-CoV infected cells, whereas a major overlap was observed between SARS-CoV-2 and MERS-CoV-2 with 13 IFN-signaling related proteins differentially regulated (Figure 6B and Figure S6). SARS-CoV-2 and SARS-CoV showed only STAT1 and EIF4A2 to be commonly upregulated (Figure 6A). The differential log2-fold change in MERS-CoV 24hpi and SARS-CoV-2 48hpi are represented as volcano plots (Figure 6C and 6D respectively). Of the 13 commonly regulated proteins between SARS-CoV-2 and MERS-CoV ISG15, IFIT1, EIF2AK, NUP54, NUP93 and SEH1L were upregulated in both, JAK1 and IFI35 were downregulated in both, while PIAS1 was upregulated in SARS-CoV2 and downregulated in MERS-CoV and nuclear receptors like KPNA1, KPNA2 and RAE1 were downregulated in SARS-CoV-2 and up-regulated in MERS-CoV. The individual protein network showing the differentially regulated proteins in the IFN-signaling pathway is shown in Figure S7. Cumulatively, this data shows distinct pattern of regulation of type-I IFN response in these three viruses.

**Figure 6.**
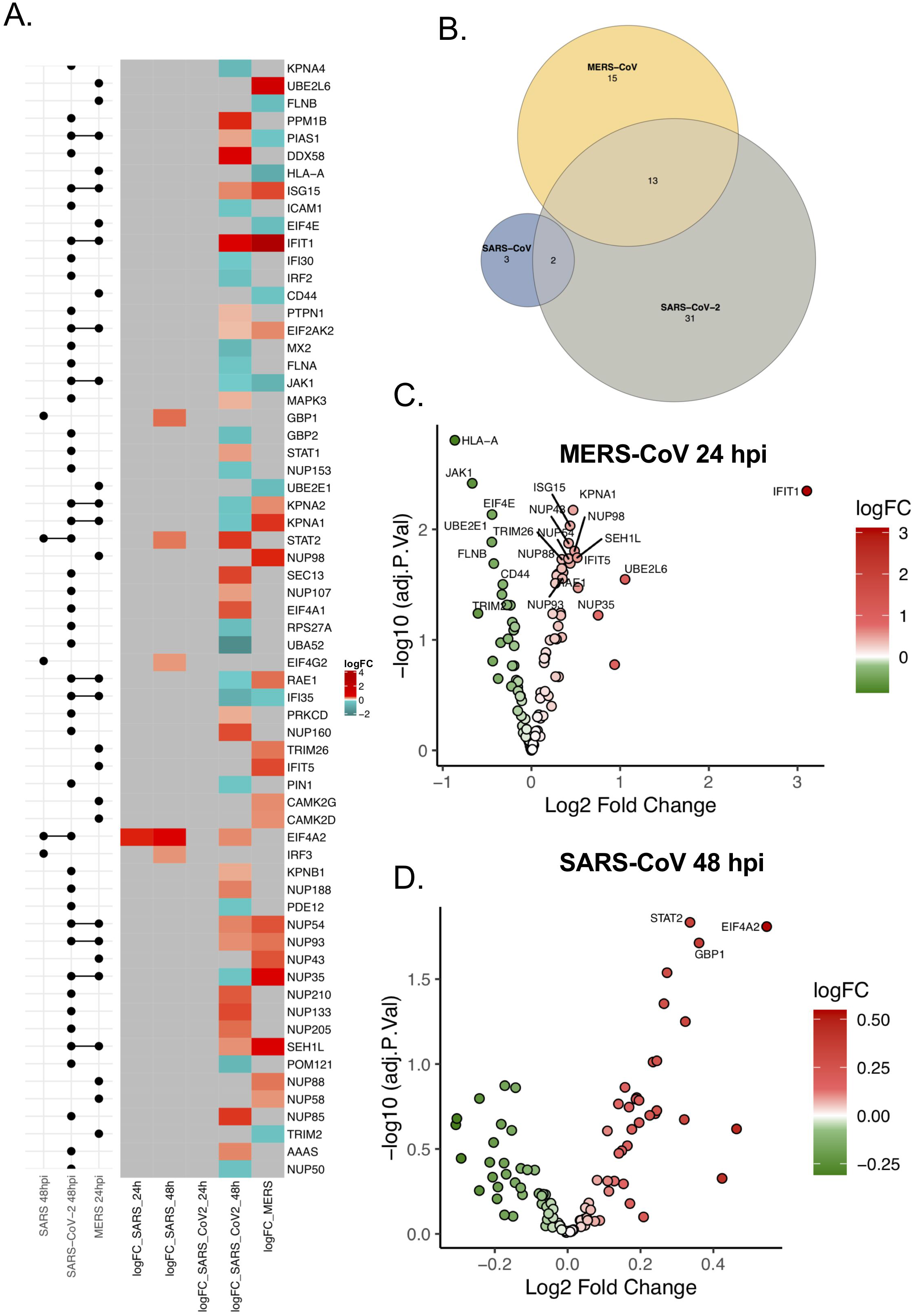
Differential regulation of IFN response by SARS-CoV, SARS-CoV-2 and MERS-CoV. A) Heatmap of log fold changes of proteins associated with IFN-signaling during SARS-CoV, SARS-CoV-2 and MERS-CoV infections. LogFC between mock and virus-infected Huh7 cells at 24hpi and 48hpi (right panel). Log fold changes associated with non-significant proteins are represented in grey. Log fold changes associated with significant differential expressed downregulated proteins are indicated in turquoise and upregulated proteins in red. The left panel of the graph shows the matrix that indicates intersects between comparisons (Mock/uninfected) using horizontal bars. B) Venn diagram illustrating the overlap between the regulated proteins belonging to the IFN-signaling by the three viruses. C) C) Volcano Plots of IFN-signaling associated proteins with differential abundance between Mock and MERS-CoV infected cells at 24hpi. D) D) Volcano Plots of IFN-signaling associated proteins with differential abundance between Mock and SARS-CoV infected cells at 48hpi.

## Discussion

The impact of the viral infection is most often dictated by the host innate immune responses and the ability of the virus to regulate these antiviral responses. Type-I interferon (IFN-I) response is one of the earliest antiviral innate immune responses following virus infection. In the present study using a proteomics-based approach, we show that SARS-CoV-2 infection induces a dysregulated IFN-I signaling in a delayed manner in Huh7 cells. Furthermore, comparison between SARS-CoV-2, SARS-CoV and MERS-CoV revealed a differential regulatory signature of interferon-related proteins.

In case of RNA viruses IFN-I response is usually initiated by recognition of the viral RNA by pattern recognition receptors (PRRs) like RIG-I and MDA-5 ^7,26^. Activation of RIG-I and MDA-5 leads to signaling cascades that are tightly controlled by post-translational modifications like ubiquitination, ISGylation and phosphorylation. The Phosphorylation of IRF3 a downstream effector of this cascade and a transcription factor leads to its dimerization and entry into the nucleus where it binds to IFN-β gene regulatory elements leading to production and release of IFN-β ^27^. The released IFN-β can further bind to interferon-α/β receptors (IFNARs) in bystander cells and initiate JAK-STAT signaling cascade, where STAT1 and STAT2 are phosphorylated and forms either a homo-dimer or hetero-dimer, which drives transcription of several interferon stimulatory genes (ISGs) upon translocation to the nucleus and ^28^. In our proteomics data we observed several components of this signaling pathway to be dysregulated and the proteomic changes are delayed by 48hrs after infection in Huh7 cells (Figure 1). In concordance with the delay in induction of ISGs, we have observed that SARS-CoV-2 can inhibit IFN-β production (Figure 3F and 3H). However, it needs to be noted that while SARS-CoV-2 induced several ISGs, many of them like MX2, GBP2, IFI30, IFI35 etc. were suppressed. Most interestingly even though several ISGs were induced, JAK1 levels were suppressed, which can make the infected cells resistant towards IFN-treatment at later stages ^29^. Other than the ISG’s several nuclear transporter complexes were also differentially modulated.

Like any other pathogenic virus, SARS-CoV-2 has developed mechanisms to suppress IFN-response. For example, by SARS-CoV-2 proteins interacting with various components of the host innate immune responses ^30^. ORF6, nsp6, nsp13, nsp1 and M proteins has been shown to inhibit IFN-I signaling pathway at different levels ^10,22,31^. On the other hand, several SARS-CoV-2 proteins like nsp2 and S proteins were found to stimulate IFN response ^22^. Thus, SARS-CoV-2 has the ability to modulate the IFN signaling in both positive and negative ways. This is represented in our findings of both increased expression and suppression of many ISGs in the infected Huh7. Not only ISGs, but also the expression of several nuclear pore complexes (involved in STAT translocation to the nucleus and subsequent ISRE-dependent gene activation) was altered in our infection model. Among the nuclear transporters Nup98 is the most studied with respect to SARS-CoV-2 infection as the ORF-6 protein interacts with it and blocks the translocation of STAT-1 to the nucleus to inhibit ISGs ^31^. However, we did not observe any change in Nup98 expression levels. Interestingly we detected another family of nuclear transporter KPNA1, KPNA2 and KPNA4 to be significantly decreased at the later time point of infection (Figure 1). KPNA1 forms a complex with pSTAT1 and aids in its translocation to the nucleus ^32^ and thus serves a major purpose in transcription of ISGs. Reduced expression of KPNA’s could result in insufficient nuclear translocation of p-STATs and thus suppress expression of many of the ISGs. Several viruses, like foot-and-mouth disease virus (FMDV), can degrade KPNA1 to block ISGs by their 3C-like protease activity ^33^ that is also encoded in ORF1a of coronaviruses and was detected in proteomics ^19^. SARS-CoV-2 also encodes another protease, papain-like protease (PLpro) that has de-ubiquitinase and de-ISGylase activity. PLpro can hydrolyze ubiquitin and ISG15 conjugation and has been implicated in SARS-CoV-2 immune evasion strategies. PLpro was also reported to be a stronger de-ISGylase than a de-ubiquitinase compared to SARS-CoV and MERS-CoV PLpro ^11^. Based on our observation of a dose-dependent decrease in conjugated-ISG15 levels at 24hpi and thereafter increase at later stages (48h) of infection, it is tempting to speculate that PLpro may play a significant role in early infection, that requires further validation.

IFN-I pathway is of significance in SARS-CoV-2 pathogenesis because IFN-I has been considered as a major treatment choice ^34,35^. Furthermore, in severe COVID-19 patients and Ferret models in spite of a cytokine storm and induction of ISGs, a very low-level of circulating IFN-I was noted ^36–38^. This was particularly interesting since in our infection model we did not observe any significant transcriptional activation of IFN-β in qPCR, despite observing changes in the levels of proteins related to RIG-I signaling and ISGs (Figure 1). A recurrent observation was the absence of correlation between transcript levels and protein levels, as both in qPCR and in transcriptomics data we did not observe any significant changes in ISG15, IFIT1, MX2, DDX58 mRNAs between the mock infected and SARS-CoV-2 infected after 48h (Figure 2 and Figure 4). In our previous paper, we observed significant changes in the level of global transcripts only after 72 h of infection. In concordance with earlier studies ^17,31,39^, we observed that IFN-pre-sensitized cells were more resistant to SARS-CoV-2, but IFN-treatment following infection did not alter the susceptibility of the cells. These results suggest that IFN-treatment may be effective in curbing SARS-CoV-2 infection, which was observed in a phase-2 trial with nebulized IFN-β-1a showing better recovery in COVID-19 patients ^40^. However, it needs to be used with caution since it may be effective when administered very early during the disease, but the virus can be resistant to the late administration of IFN or late induction of ISGs following established virus replication that can further contribute to the pathology. Therefore, we are postulating that people with naturally high level of IFN may have the potency to control the virus early and moving the balance towards better disease outcome.

Older people are at a higher risk of COVID-19 with increased risk of severe disease ^25^ that could be attributed to the cellular senescence associated with age. Senescent cells secrete a plethora of mediators (senescence associated secretory phenotype, SASP), many with pro-inflammatory activity and show highly dysregulated immune response ^41^. There are contradictory evidences of both inhibition and enhancement of viral replication in senescent cells ^42^. Senescent human bronchial epithelial cells were observed to be more susceptible to influenza A virus ^43^. However, presently no data is available on SARS-CoV-2 susceptibility in senescent cells and the type of response it can exhibit.

In our Huh7 senescent cell model even though there was a significant increase in IFN-response compared to healthy cells, the virus production was significantly increased (Figure 5), suggesting that the virus is able to escape the antiviral response in senescent cells. In particular among the ISGs tested we observed a significant suppression of IFIT1. However, this effect may be cell-type dependent. For instance, Caco2 cells showed more resistance to etoposide with a very low-level induction of senescence as represented by p21. However, we observed an inhibition of viral replication with visible upregulation of IFIT1. This indicates, IFIT1 to be an important antiviral-factor that needs further attention. Also, the differences observed among the two cell lines underscores the drawback of studying a single cell line (Huh7 in this case) as it may not be reflective of other cell populations where there could be differential regulation of IFN-response.

SARS-CoV-2 shows a higher level of susceptibility to IFN-treatment in comparison to SARS-CoV ^17^ and its sensitivity to IFN-I pretreatment is shared by MERS-CoV ^10,17,44,45^. In the Huh7 infection model, we have observed the MERS-CoV to be highly cytopathic, a delayed cytopathic effect in SARS-CoV and no cytopathic effect with SARS-CoV-2 infection at the same infective dose. This points towards a differential regulation of immune-signaling pathways by these viruses. Using proteomics, we attempted to delineate the immunological features of the cells during infection with these three viruses. We were restricted with our analysis of MERS-CoV to 24hpi and we observed a large number of proteins expression to be significantly altered when compared to the mock. While in case of SARS-CoV and SARS-CoV-2 the major changes were observed at 48hpi. While we observed a variety of cellular processes to be commonly regulated by these viruses (Supplementary Fig S5), we focused our analysis to IFN-I signaling. All the three viruses had unique signatures in induction of IFN-response in Huh7 cells, with very limited overlap among them. While SARS-CoV-2 and MERS-CoV had many similar signatures, SARS-CoV showed very little induction of ISG’s and there was no similarity to MERS-CoV at all (Figure 6). This probably explains the resistance to IFN-treatment observed in SARS-CoV in other studies ^17^, as it may have a stronger mechanism to inhibit IFN-I response. SARS-CoV-2 and MERS-CoV had 13 common proteins that were significantly altered. However, while the nuclear transporter complex proteins KPNA1, KPNA2, RAE1 were suppressed in SARS-CoV-2 infected cells, they were upregulated in MERS-CoV infected cells. Earlier we have discussed the possible role of 3C-like protease encoded in ORF3a in degradation of KPNA isoforms. The absence of visible detection of ORF1a or 3CL-pro peptides in MERS-CoV infected cells further strengthens the role of these viral proteins in regulation of transport of cellular transcription factors to the nucleus.

To conclude, our findings provide a better understanding of the regulation of cellular interferon response during SARS-CoV-2 infection and a perspective on the use of interferons as a treatment. The proteomics findings highlight that SARS-CoV-2 related human pathogenic coronaviruses regulate the IFN-signaling differently and previous findings on SARS-CoV and MERS-CoV should not be automatically applied on SARS-CoV-2. Detailed characterization of the role of different ISGs on inhibition of SARS-CoV-2 pathogenesis may direct novel antiviral strategies.

## Materials and Methods

### Chemicals

Bovine serum albumin (BSA, A7906) were purchased from Sigma-Aldrich (USA). 10% sodium dodecyl sulfate (SDS), 0.5M ethylenediaminetetraacetic acid disodium salt dehydrate (EDTA), 5M sodium chloride (NaCl), 1M Tris base pH 7.6 and 20% Tween-20 was purchased from Karolinska Institutet substrate department (Sweden). Poly(I:C) (LMW)/LyoVec was purchased lyophilized from InvivoGen (France) and resuspended in sterile physiological water at a final concentration of 20 mg/mL. Acitretin (44707) and Etoposide (E1383) was purchased from Sigma-Aldrich (USA). Interferon-α (IFN-α) and interferon-β (IFN-β) were purchased from PBL (USA).

### Antibodies

Antibodies and their manufacturers were: rabbit anti-RIG-I clone D14G6 (1:1000; #3743), rabbit anti-MDA5 clone D74E4 (1:1000; #5321) from Cell-Signaling Technologies (Danvers, MA, USA), mouse anti-ISG15 (1:1000, sc-166755) from Santa-Cruz Biotechnology (santa Cruz, CA, USA), recombinant Anti-GAPDH clone EPR16891(1:10000, Ab181602) and rabbit anti TRIM25 clone EPR7315 (1:2000; ab167154) from Abcam (Cambridge, MA, USA).

### Cell lines and virus

The human hepatocyte-derived cellular carcinoma Huh7 cell line was obtained from Marburg Virology Lab, Germany and Caco2 was obtained from CLS cell line services, GmbH, Germany (#300137). The cell lines were maintained in Dulbecco’s modified Eagle medium (DMEM, ThermoFisher, USA) supplemented with 10% fetal bovine serum (FBS, ThermoFisher, USA) and 20 units/mL penicillin combined with 20 μg/mL streptomycin (Sigma, USA). Cells were cultured in 5% CO_2_ at 37°C.

The SARS-CoV-2 virus was isolated from a nasopharyngeal sample of a patient in Sweden and the isolated virus was confirmed as SARS-CoV-2 by sequencing (Genbank accession number MT093571) and titrated as described elsewhere ^19^.

### RIG-I agonist and Interferon treatment

Huh7 cells were seeded in 24-well plates (6×10^4^ cells/well) in DMEM supplemented with 10% heat-inactivated FBS; and after 24 h the cells were treated with poly I:C (10 μg/mL), acitretin (25 μM), IFN-β (100 IU) and IFN-α (5000 IU) in DMEM supplemented with 5% heat-inactivated FBS for 16 h before infection. Then, pre-treated and non-treated cells were either cultured in DMEM with 5% FBS (uninfected control) or infected with SARS-CoV-2 at a multiplicity of infection (MOI) of 0.1 added in a total volume of 0.5 mL. After one hour of incubation (37°C, 5% CO_2_) the inoculum was removed, and medium only was added to pre-treated and uninfected cells, while medium with the compounds dilutions was added for cell treatment post-infection.

### Etopside treatment

Huh7 cells were seeded in 6-well plates in DMEM supplemented with 10% heat-inactivated FBS. Cells were either treated with 0.5 μM of etoposide or left untreated. The etoposide-supplemented medium or the normal medium was replenished after 3 days. Following 6 days of etoposide treatment the cells were left in normal medium for 1 day and then they were split into 12-well plate at a seeding density of 25,000 cells/well in 1 mL of normal-medium. Twenty-four hours post-seeding the cells were either mock infected or SARS-CoV-2 infected (MOI 0.1) in triplicate for 1hr followed by replenishing the medium with DMEM containing 5% FBS. The supernatant and the cells were harvested 24 h after infection to determine the virus production and the mRNA levels of the proteins of interest.

The cell culture supernatant was collected 24 h post-infection and stored for viral load quantification, while cells were collected by adding Trizol™ (ThermoFisher Scientific, US) directly to the wells. RNA was extracted from SARS-CoV-2 infected and uninfected Huh7 cells using the Direct-zol™ RNA Miniprep (Zymo Research, USA).

### Immunoblots

Following 24hpi and 48hpi infection with different doses of SARS-CoV-2, the cells were lysed in 2% SDS lysis buffer (50 mM Tris-Cl pH 7.4, 150 mM NaCl, 1 mM EDTA, 2% SDS, freshly supplemented with 1 mM dithiothreitol (DTT), 1x protease inhibitor cocktail and 1x phosphatase inhibitor cocktail) followed by boiling at 95°C for 10 minutes to inactivate the virus. The protein concentration was evaluated by DC Protein Assay from Bio-Rad (USA). Evaluation of protein expression was performed by running 20 μg of total protein lysate on NuPage Bis Tris 4%-12% gels (Invitrogen, USA). Proteins were transferred using iBlot dry transfer system (Invitrogen, USA) and blocked for one hour using 5% milk or BSA in 0.1% PBS-t (0.1% Tween-20). Subsequent antibody incubation was performed at 4°C overnight or for one hour at room temperature using Dako polyclonal goat anti-rabbit or anti-mouse immunoglobulins/HRP (Agilent Technologies, USA). Membranes were washed using 0.1% TBS-T and proteins were detected using ECL or ECL Select (GE Healthcare, USA) on ChemiDoc XRS+ System (Bio-Rad Laboratories, USA). The Western blot analysis was performed by using antibodies targeting RIG-I, MDA-5, TRIM25, ISG15, GAPDH.

### Quantitative RT-PCR

Viral RNA was quantified from cell supernatant as a confirmation of the infection by Takara PrimeDirect probe, RT-qPCR mix (Takara Bio Inc., Japan), with primers and probe specific for the SARS-CoV-2 E gene, as previously reported ^46^. The Primers and probes used were E_Sarbeco_F1: 5’-ACAGGTACGTTAATAGTTAATAGCGT-3’, E_Sarbeco_R2: 5’-ATATTGCAGCAGTACGCACACA-3’ and Probe: [FAM] ACACTAGCCATCCTTACTGCGCTTCG [BBQ650]. RT-qPCR was performed using 400 nM of primers and 200 nM of probe with cycling conditions: initial denaturation at 90°C for 3 min, reverse transcription at 60°C for 5 min, followed by 45 cycles of 95°C for 5 s and 58°C for 30 s. Messenger RNA (mRNA) expression of a few ISGs transcripts and human GAPDH was measured by qRT-PCR. The sequences of the qPCR primers are listed in supporting information (Table S1). Total RNA was extracted using Direct-zol™ RNA miniprep (Zymo Research, USA) and RNA concentration was assessed using a spectrophotometer (NanoDrop UV Visible Spectrophotometer, Thermofisher, USA). Reverse transcription was performed using a high capacity reverse transcription kit (Applied Biosystems, USA) or SuperScript vilo cDNA synthesis kit (Thermofisher, USA) for 10 min at 25°C, followed by 37°C for 120 min and 85°C for 5 min. Quantitative RT-PCR assays were setup using the Power SYBR Green PCR Master Mix (Applied Biosystems, UK) using 250 nM of primer pairs with cycling conditions: initial denaturation 95°C 10 min, followed by 40 cycles of 95°C for 15 s, 60°C for 1 min. Melting curves were run by incubating the reaction mixtures at 95°C for 15 s, 60°C for 20 s, 95°C for 15 s, ramping from 60°C to 95°C in 1°C/s. The values were normalized to endogenous GAPDH.

Fold change was calculated as: Fold Change = 2-Δ(ΔCt) where ΔCt = Ct target—Ct housekeeping and Δ(ΔCT) = ΔCt infected - ΔCt mock-infected/untreated, according to the Minimum Information for Publication of Quantitative Real-Time PCR Experiments (MIQE) guidelines.

### Quantitative proteomics analysis

Proteomics workflow was performed similarly as we described previously ^19^. Briefly, proteins were extracted with SDS-based buffer, digestion was performed on S-Trap micro columns (Protifi, Huntington, NY) and resulting peptides were labeled with isobaric TMTpro™ reagents. Labeled peptides were fractionated by high pH (HpH) reversed-phase chromatography, and each fraction was analyzed on an Ultimate 3000 UHPLC (ThermoFisher Scientific) in a 120 min linear gradient. Data were acquired on a Orbitrap Fusion Lumos™ tribrid mass spectrometer (ThermoFisher Scientific) in data dependent acquisition (DDA) mode, isolating precursors in 2 s cycle time with 120,000 mass resolution in the mass range of 375 – 1500 *m/z*, maximum injection time (IT) of 50 ms and dynamic exclusion of 45 s; precursor isolation width of 0.7 Th with high collision energy (HCD) of 34%, resolution of 30,000 and maximum IT of 54 ms.

Proteins were searched against both SwissProt human and SARS-CoV/SARS-CoV2 databases using the search engine Mascot Server v2.5.1 (MatrixScience Ltd, UK) in Proteome Discoverer v2.4 (ThermoFisher Scientific) software allowing up to two missed cleavages. Oxidation of methionine, deamidation of asparagine and glutamine, TMTpro modification of lysine and N-termini were set as variable modifications; while carbamidomethylation of cysteine was used as fixed modification. The false discovery rate (FDR) was set to 1%. The raw mass spectrometric data was deposited to the ProteomeXhanger Consortium (http://proteomecentral.proteomexchange.org) via the PRIDE partner repository with the dataset identifier PXD023450.

### Statistical analysis

Statistical analyses for proteomics and transcriptomics were performed in R package LIMMA. All other statistical calculations were performed in GraphPad Prism (Version 8.0.0) using unpaired t-test. Significance values are indicated in the figures and figure legends. p*<0.05, **<0.01, ***<0.001 and ****<0.0001.

### Bioinformatics analysis

Proteo-transcriptomics data of SARS-CoV-2 infected (1 MOI) Huh7 cells to identify the temporal pattern changes resulting from infection were re-analyzed ^19^. Huh7 cells infected with SARS-CoV-2 at MOI 1 were collected at 24, 48, and 72 h in triplicates. Differential abundance analysis was performed using R package LIMMA between mock infected and respectively 24 and 48hpi in transcriptomics and proteomics data. Pairwise comparisons were extracted and Benjamini-Hochberg (BH) adjustment was applied on p-values. Genes with adjusted p-values <0.05 were selected. Three manually curated libraries based on interferon-regulated genes were created based on reactome terms “Antiviral mechanism by IFN−stimulated genes”, “Interferon gamma signaling” and “Interferon alpha/beta signaling” (https://reactome.org/). Each library had respectively 89, 172 and 138 genes. The total number of interferon-regulated genes excluding overlap between libraries is 205. Among this set, 97 proteins and 144 genes were detected in the data. Proteins and transcripts profiles were represented as a heatmap using R ComplexHeatmap function. 48 proteins and 8 genes were significantly changing between mock and 48hpi. Interferon-regulated genes and proteins from differential abundance analysis were extracted and represented as volcano plot using ggplot2. Significant proteins (proteomics data, LIMMA, FDR < 0.05) were represented as a network with Cytoscape ver 3.6.1. For each node, fold changes were added to the network template file. Protein-protein interactions were retrieved from STRING Db (v5.0) (https://string-db.org/). Interactions were filtered on confidence score with minimum interaction of 0,700. Only interactions from databases and experiences were conserved. Genes associated with type I interferon identified in proteomics data were represented as dot plots using ggplot2.

Huh7 cells infected were collected at 24 and 48hpi for SARS-CoV and 24hpi for MERS-CoV. Mock infected cells were collected at similar time points. Proteomics raw data was first filtered for empty rows and quantile normalized with R package NormalizerDE. Histogram was used to display the distribution of data and assess that the distribution follows a normal law. Principal component analysis was performed using ggplot2. Viral protein abundances were retrieved and baseline subtraction (Infected-Mock) was performed for each time point and represented using barplots made with ggplot2. In order to identify proteins changing after infection, differential abundance analysis was performed using R package LIMMA between Mock and infected cells as described from Huh7 cells with SARS-CoV-2 infection. As described previously, results were filtered for interferon related libraries. 99 interferon-related proteins were detected for SARS-CoV, only 1 significant at 24 h and 5 at 48 h. For SARS-CoV and MERS-CoV, 96 interferon proteins were detected and 28 were differentially expressed. for Results from each comparison were retrieved and represented as volcano plot using ggplot2, Venn diagram using interactivenn (http://www.interactivenn.net/) and heatmap of fold changes using R package complexHeatmap. Significant proteins identified in Huh7 infected with SARS-CoV-2, SARS-CoV and MERS-CoV were extracted from proteomic data and represented as a network.

## Supplementary Figure Legends

**Supplementary Figure S1:**
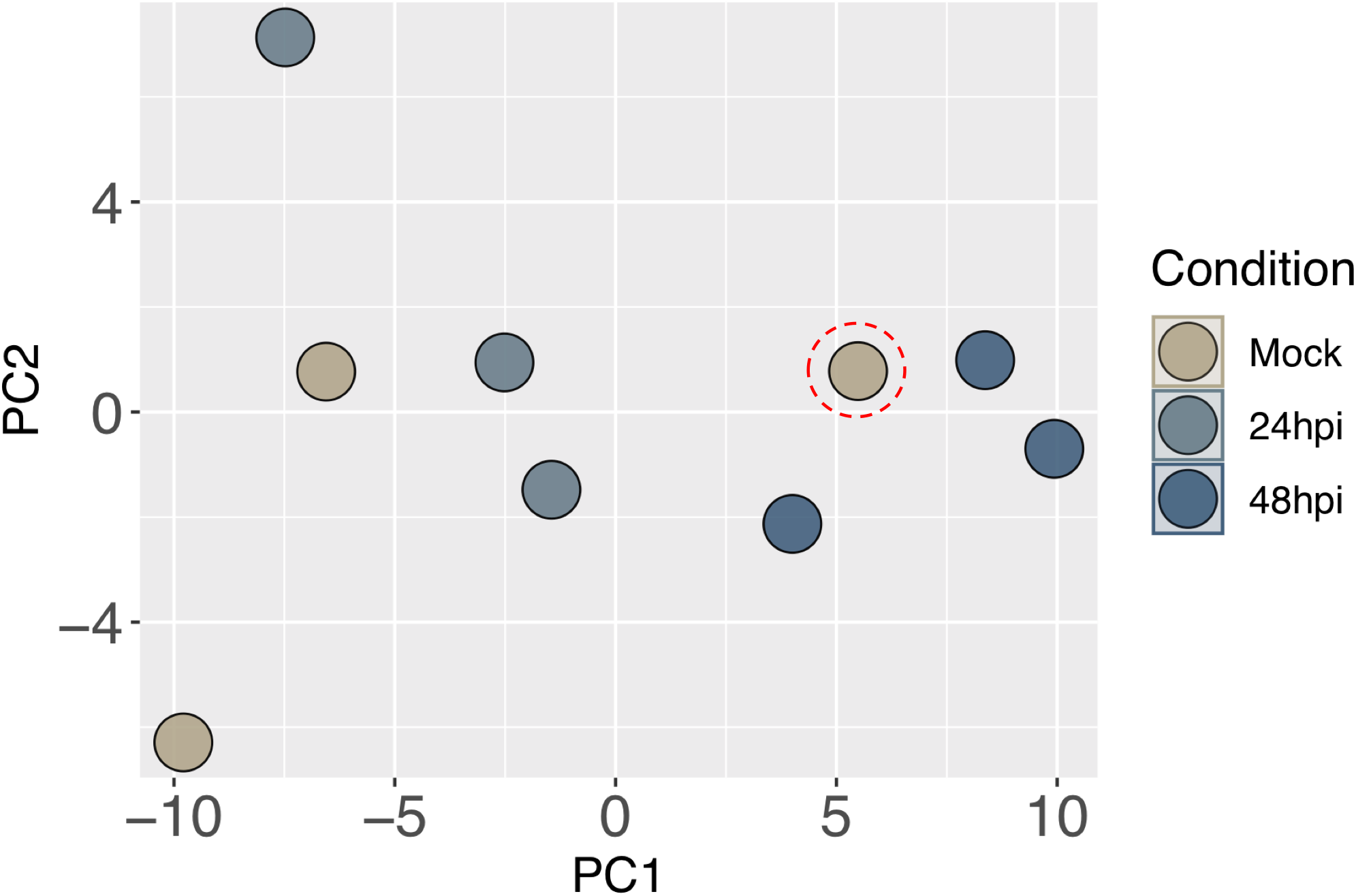
Principal component Analysis (PCA) plot of Mock infected (24 h) and SARS-CoV-2 infected (24 h and 48 h) proteomics data. One mock sample was an outlier as indicated by red dotted circle.

**Supplementary Figure S2:**
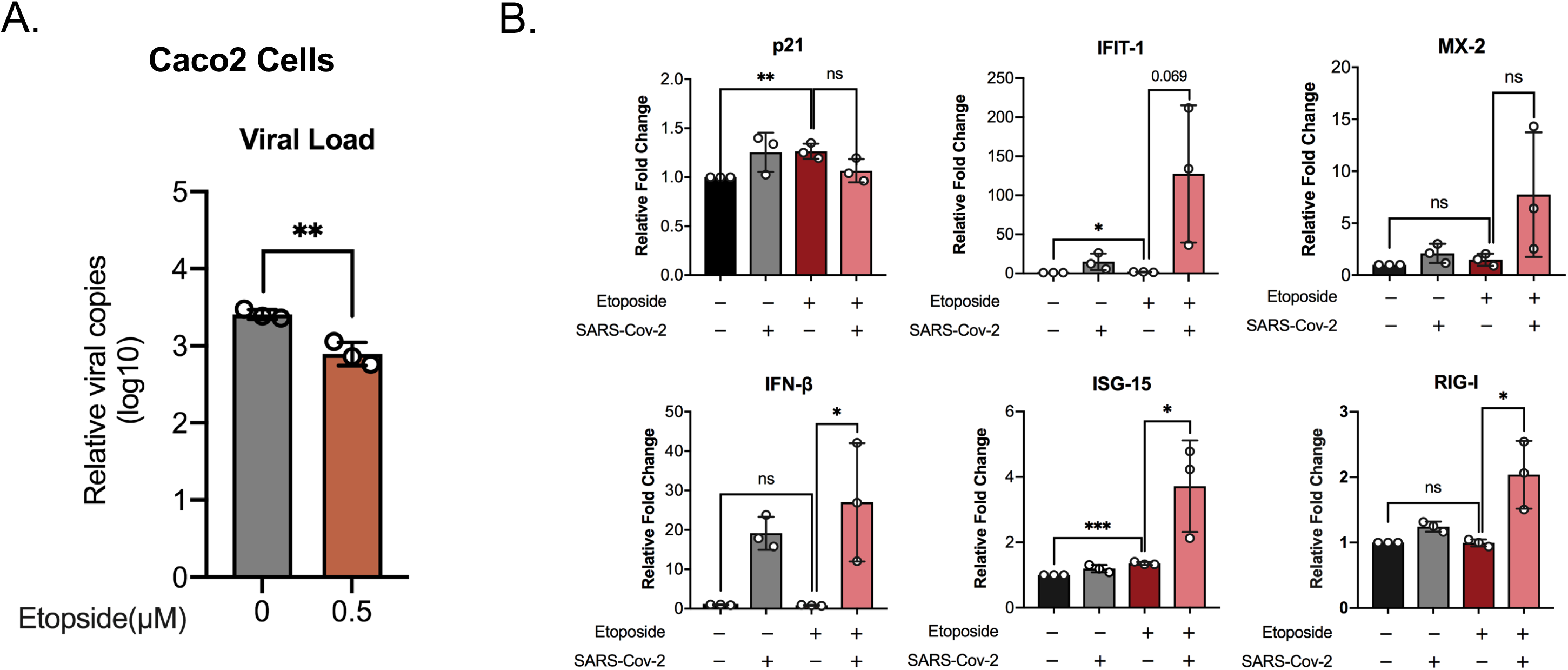
SARS-CoV-2 susceptibility in Etoposide treated Caco2 cells. Caco2 cells were treated with 0.5 μM of etoposide for 6 days followed by 2 days regular DMEM with 10% FBS. The cells were either mock infected or infected with SARS-CoV-2 at MOI of After 24 h the cell-culture supernatant and cells were harvested to determine the virus production and the transcript levels of the indicated genes respectively. The experiments were performed in technical triplicate and the mean ± SD values is shown. An unpaired t-test was used to determine p-values (* <0.05, **< 0.01, ****< 0.001) A) The virus production in Etoposide treated Caco2 cells. B) The levels of specific mRNAs were quantified by qRT-PCR. The results are shown as fold change relative to non-treated cells. The mean± SD of technical triplicates are shown.

**Supplementary Figure S3:**
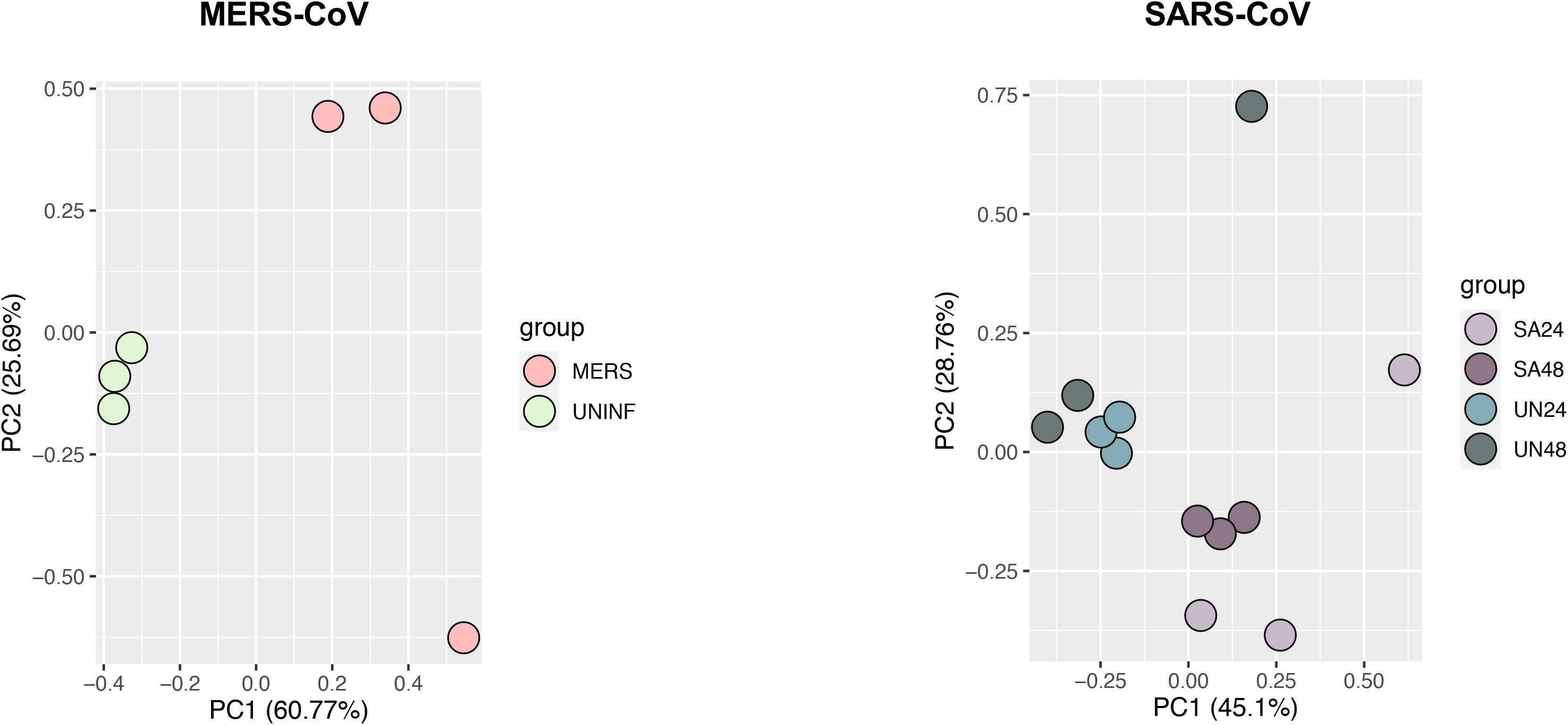
Principal component Analysis (PCA) plot of mock and MERS-CoV infected (24 h) (left panel) and of mock and SARS-CoV infected (24 h and 48 h) proteomics data (right panel).

**Supplementary Figure S4:**
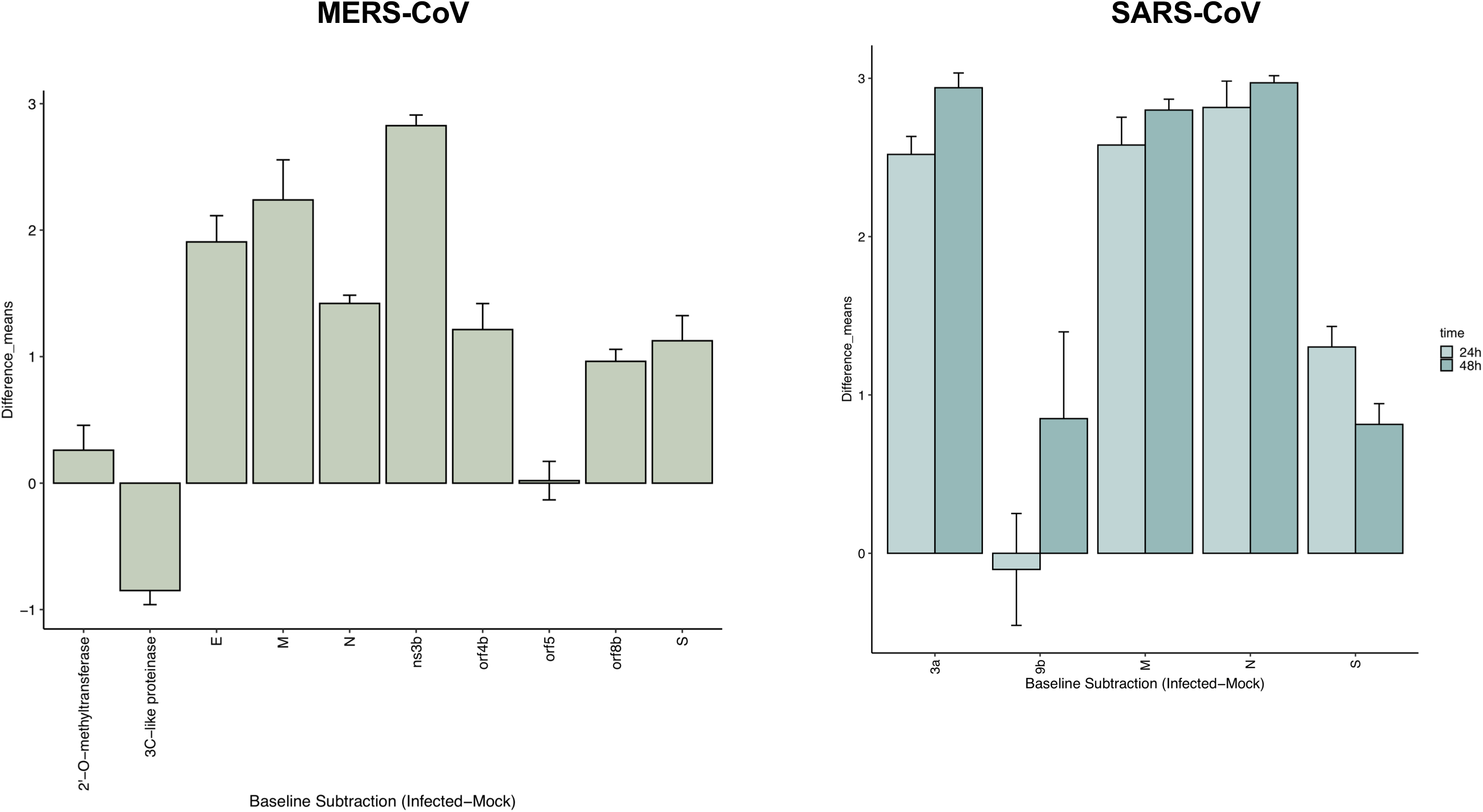
Temporal dynamics of detected viral proteins in the Huh7 cells by tandem mass tag-labelled mass spectrometry (TMT-MS). Left panel shows MERS-CoV infected cells and right panel shows SARS-CoV infected cells.

**Supplementary Figure S5:**
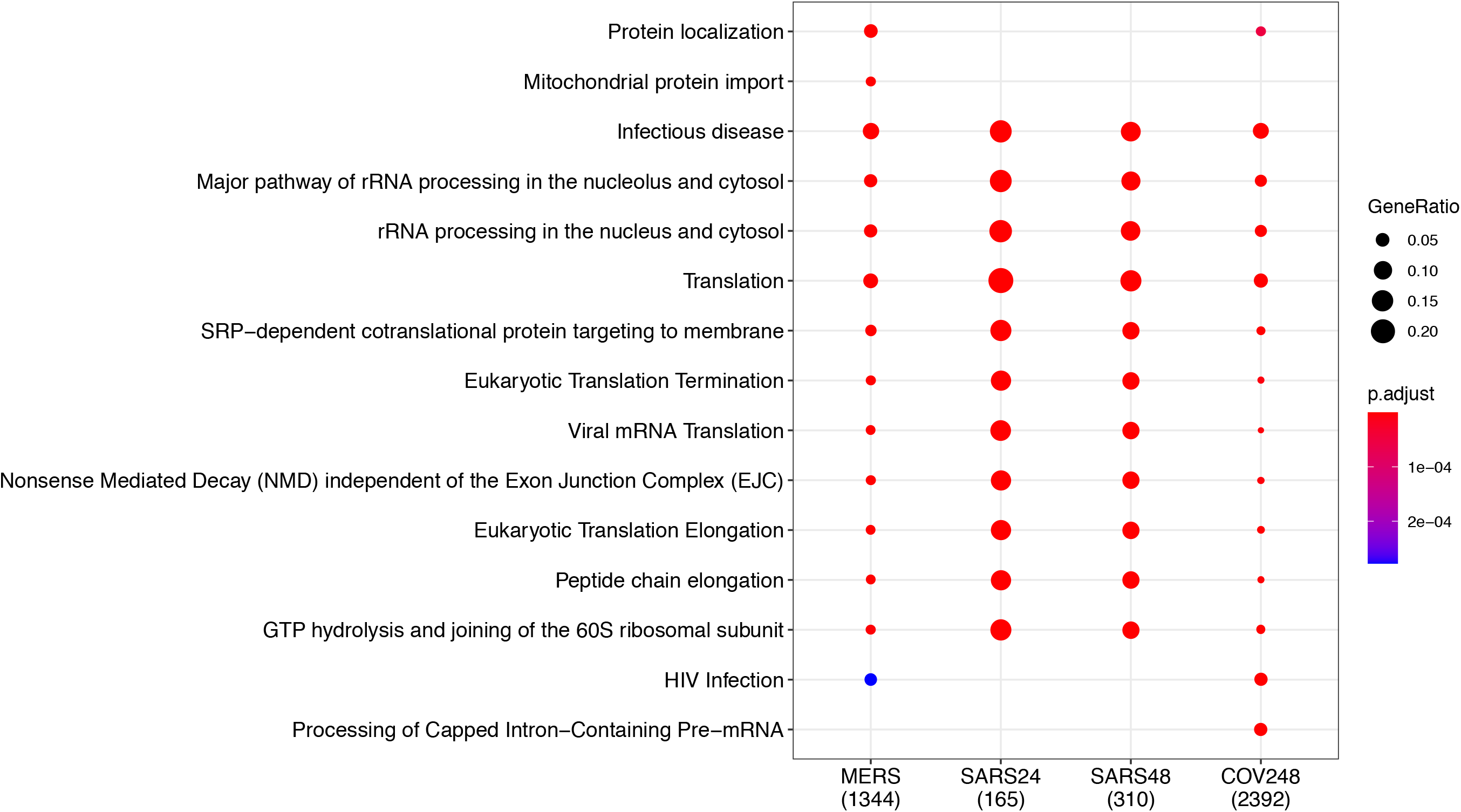
Dotplot visualization of enriched pathways. The size represents the gene ratio between enriched and total gene set. The color represents the adjusted p-value.

**Supplementary Figure S6:**
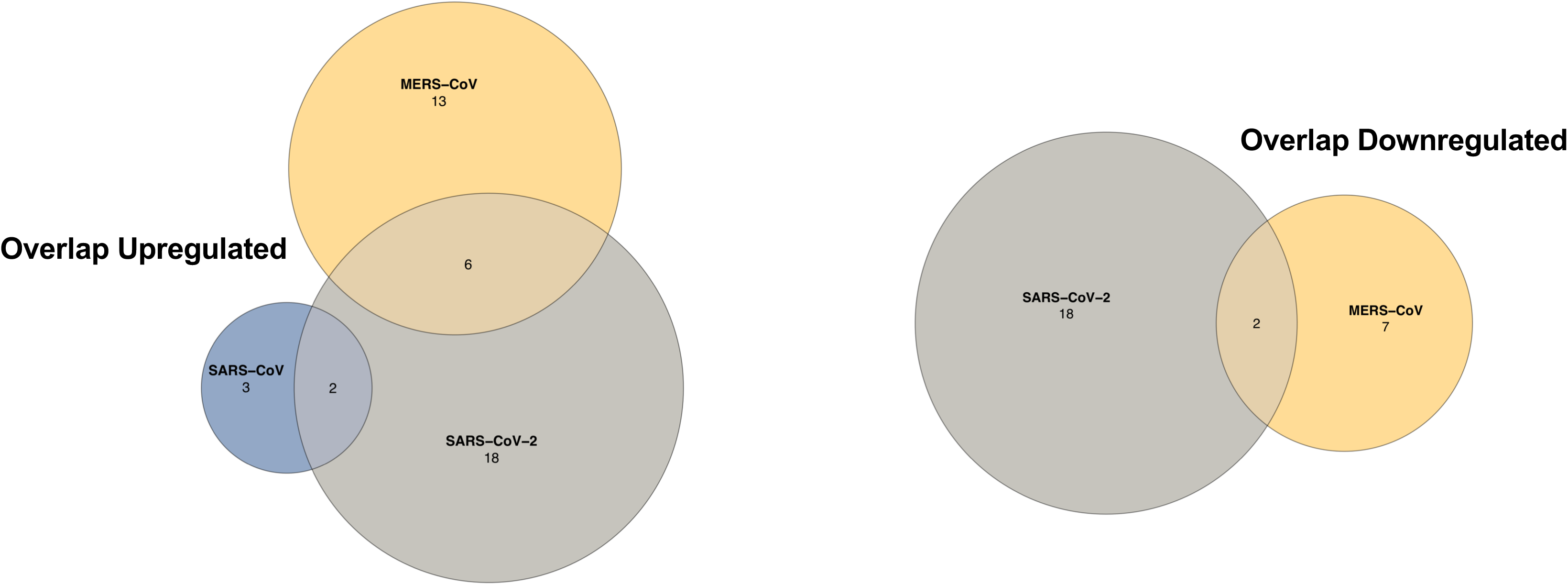
Venn diagram of IFN-signaling related proteins upregulated (left panel) and downregulated (right panel) in SARS-CoV, SARS-CoV-2 and MERS-CoV infected cells compared to mock infected cells.

**Supplementary Figure S7:**
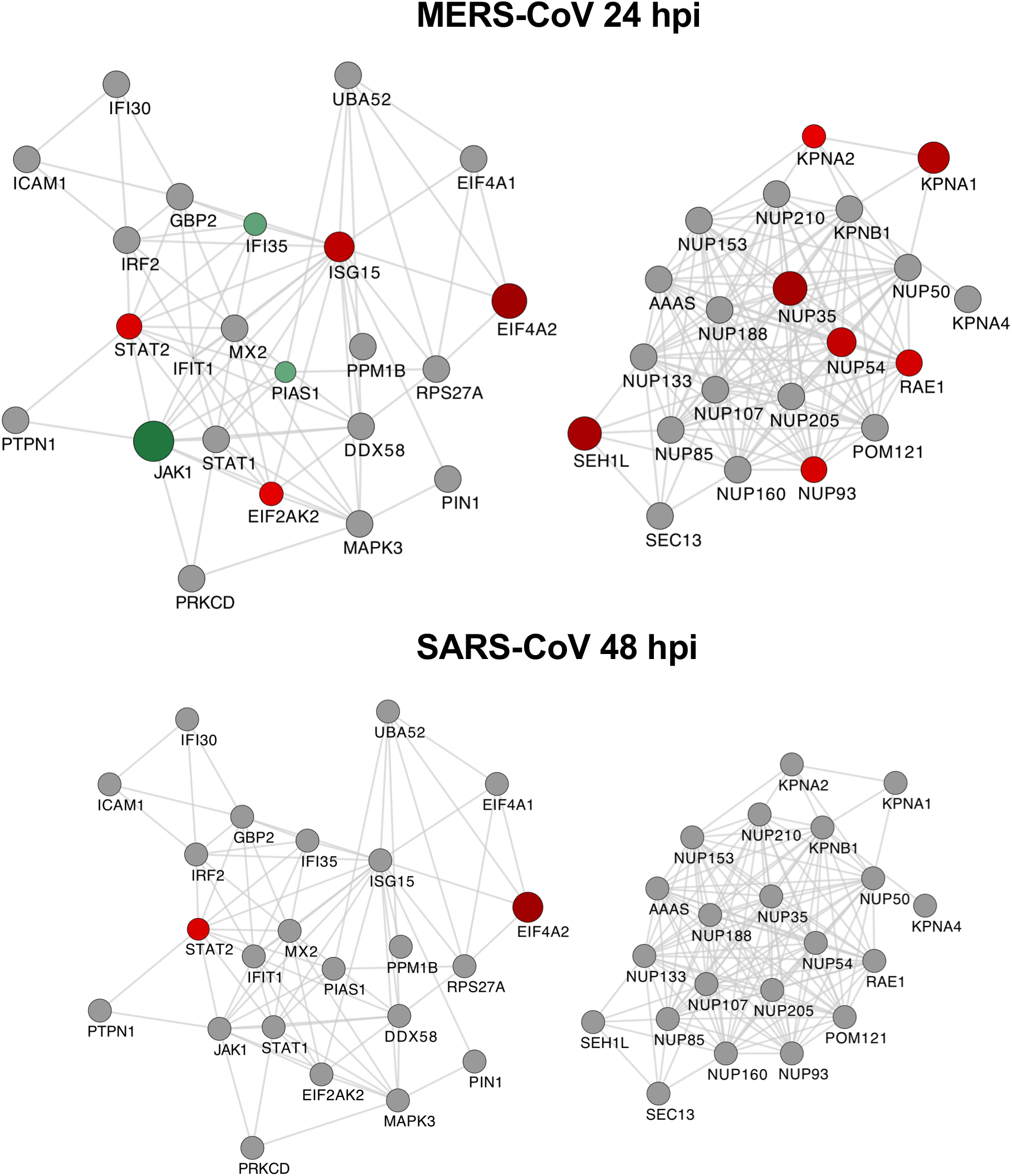
Cytoscape network of differentially abundant IFN-signaling related proteins in MERS-CoV at 24hpi (upper panel) and SARS-CoV at 48hpi (lower panel).

## Acknowledgments

The authors would like to acknowledge the support from the Proteomics Biomedicum, Karolinska Institutet for LC-MS/MS analysis. The study is funded by Swedish Research Council Grants (2017-01330) to U.N., Karolinska Institute Stiftelser och Fonder (2020-02153 to S.G. and 2020-01554 to U.N.), Åke Wibergs Stiftelse (M20-0220) to S.G., Swedish research Council (2018-05766 and 2017-03126) and Innovative Medicines Initiative 2 Joint Undertaking (JU) under grant agreement no. 101005026 to A.M. JU receives support from the European Union’s Horizon 2020 research and innovation programme and EFPIA. TF acknowledges the grant received from the Swedish Cancer Society and the Swedish Research Council.

## Data availability

The raw mass spectrometric data was deposited to the ProteomeXhanger Consortium (http://proteomecentral.proteomexchange.org) via the PRIDE partner repository with the dataset identifier PXD023450. All the bioinformatic analysis codes are available in github at https://github.com/neogilab/COVID_IFN. Additional datasets generated for this study are available on request to the corresponding author.

## Authors contribution

S.G., U.N. A.M. and T.F. conceptualized the study. E.S., X.C., K.A.S, and S.G. performed the experiments. F.M. performed the bioinformatics analysis. Á.V. and J.E.R. performed the mass-spectrometry. S.G. supervised the study. U.N. and A.M. contributed with resources. T.F. provided critical intellectual inputs. S.G. wrote the first draft of the manuscript. E.S., F.M., B.S.V., X.C., Á.V. and J.E.R. helped in writing the first draft of the manuscript. U.N. and T.F. edited the manuscript. All of the authors contributed to revising the manuscript and approved the final version of the manuscript.

## References

1 Baric, R. S. Emergence of a Highly Fit SARS-CoV-2 Variant. N Engl J Med, doi:10.1056/NEJMcibr2032888 (2020).

2 Rabaan, A. A. et al. SARS-CoV-2, SARS-CoV, and MERS-COV: A comparative overview. Infez Med 28, 174–184 (2020).

3 Chen, B. et al. Overview of lethal human coronaviruses. Signal Transduct Target Ther 5, 89, doi:10.1038/s41392-020-0190-2 (2020).

4 Lui, G. C. et al. Significantly Lower Case-fatality Ratio of Coronavirus Disease 2019 (COVID-19) than Severe Acute Respiratory Syndrome (SARS) in Hong Kong-A Territory-Wide Cohort Study. Clin Infect Dis, doi:10.1093/cid/ciaa1187 (2020).

5 Acharya, D., Liu, G. & Gack, M. U. Dysregulation of type I interferon responses in COVID-19. Nat Rev Immunol 20, 397–398, doi:10.1038/s41577-020-0346-x (2020).

6 McNab, F., Mayer-Barber, K., Sher, A., Wack, A. & O’Garra, A. Type I interferons in infectious disease. Nat Rev Immunol 15, 87–103, doi:10.1038/nri3787 (2015).

7 Kell, A. M. & Gale, M., Jr. RIG-I in RNA virus recognition. Virology 479-480, 110–121, doi:10.1016/j.virol.2015.02.017 (2015).

8 Schneider, W. M., Chevillotte, M. D. & Rice, C. M. Interferon-stimulated genes: a complex web of host defenses. Annu Rev Immunol 32, 513–545, doi:10.1146/annurev-immunol-032713-120231 (2014).

9 Kindler, E., Thiel, V. & Weber, F. Interaction of SARS and MERS Coronaviruses with the Antiviral Interferon Response. Adv Virus Res 96, 219–243, doi:10.1016/bs.aivir.2016.08.006 (2016).

10 Xia, H. et al. Evasion of Type I Interferon by SARS-CoV-2. Cell Rep 33, 108234, doi:10.1016/j.celrep.2020.108234 (2020).

11 Shin, D. et al. Papain-like protease regulates SARS-CoV-2 viral spread and innate immunity. Nature 587, 657–662, doi:10.1038/s41586-020-2601-5 (2020).

12 Sheahan, T. P. et al. Comparative therapeutic efficacy of remdesivir and combination lopinavir, ritonavir, and interferon beta against MERS-CoV. Nat Commun 11, 222, doi:10.1038/s41467-019-13940-6 (2020).

13 Sallard, E., Lescure, F. X., Yazdanpanah, Y., Mentre, F. & Peiffer-Smadja, N. Type 1 interferons as a potential treatment against COVID-19. Antiviral Res 178, 104791, doi:10.1016/j.antiviral.2020.104791 (2020).

14 Beigel, J. H. et al. Remdesivir for the Treatment of Covid-19 - Final Report. N Engl J Med 383, 1813–1826, doi:10.1056/NEJMoa2007764 (2020).

15 Channappanavar, R. et al. Dysregulated Type I Interferon and Inflammatory Monocyte-Macrophage Responses Cause Lethal Pneumonia in SARS-CoV-Infected Mice. Cell Host Microbe 19, 181–193, doi:10.1016/j.chom.2016.01.007 (2016).

16 Channappanavar, R. et al. IFN-I response timing relative to virus replication determines MERS coronavirus infection outcomes. J Clin Invest 129, 3625–3639, doi:10.1172/jci126363 (2019).

17 Lokugamage, K. G. et al. Type I Interferon Susceptibility Distinguishes SARS-CoV-2 from SARS-CoV. J Virol 94, doi:10.1128/jvi.01410-20 (2020).

18 Sun, J. et al. Comparative Transcriptome Analysis Reveals the Intensive Early Stage Responses of Host Cells to SARS-CoV-2 Infection. Front Microbiol 11, 593857, doi:10.3389/fmicb.2020.593857 (2020).

19 Appelberg, S. et al. Dysregulation in Akt/mTOR/HIF-1 signaling identified by proteo-transcriptomics of SARS-CoV-2 infected cells. Emerg Microbes Infect 9, 1748–1760, doi:10.1080/22221751.2020.1799723 (2020).

20 Tiwari, R. et al. In silico and in vitro studies reveal complement system drives coagulation cascade in SARS-CoV-2 pathogenesis. Comput Struct Biotechnol J 18, 3734–3744, doi:10.1016/j.csbj.2020.11.005 (2020).

21 Elisa Saccon, S. K., Beatriz Sá Vinhas, Siddappa N. Byrareddy, Ali Mirazimi, Ujjwal Neogi, Soham Gupta. Replication dynamics and cytotoxicity of SARS-CoV-2 Swedish isolate in commonly used laboratory cell lines. bioRxiv, doi:10.1101/2020.08.28.271684 (2020).

22 Lei, X. et al. Activation and evasion of type I interferon responses by SARS-CoV-2. Nat Commun 11, 3810, doi:10.1038/s41467-020-17665-9 (2020).

23 Dzimianski, J. V., Scholte, F. E. M., Bergeron, É. & Pegan, S. D. ISG15: It’s Complicated. J Mol Biol 431, 4203–4216, doi:10.1016/j.jmb.2019.03.013 (2019).

24 Perng, Y. C. & Lenschow, D. J. ISG15 in antiviral immunity and beyond. Nat Rev Microbiol 16, 423–439, doi:10.1038/s41579-018-0020-5 (2018).

25 Wang, D. et al. Clinical Characteristics of 138 Hospitalized Patients With 2019 Novel Coronavirus-Infected Pneumonia in Wuhan, China. JAMA 323, 1061–1069, doi:10.1001/jama.2020.1585 (2020).

26 Loo, Y. M. & Gale, M., Jr. Immune signaling by RIG-I-like receptors. Immunity 34, 680–692, doi:10.1016/j.immuni.2011.05.003 (2011).

27 Gupta, S. et al. 14-3-3 scaffold proteins mediate the inactivation of trim25 and inhibition of the type I interferon response by herpesvirus deconjugases. PLoS Pathog 15, e1008146, doi:10.1371/journal.ppat.1008146 (2019).

28 Ivashkiv, L. B. & Donlin, L. T. Regulation of type I interferon responses. Nat Rev Immunol 14, 36–49, doi:10.1038/nri3581 (2014).

29 Hazari, S. et al. Reduced expression of Jak-1 and Tyk-2 proteins leads to interferon resistance in hepatitis C virus replicon. Virol J 4, 89, doi:10.1186/1743-422x-4-89 (2007).

30 Gordon, D. E. et al. A SARS-CoV-2 protein interaction map reveals targets for drug repurposing. Nature 583, 459–468, doi:10.1038/s41586-020-2286-9 (2020).

31 Miorin, L. et al. SARS-CoV-2 Orf6 hijacks Nup98 to block STAT nuclear import and antagonize interferon signaling. Proc Natl Acad Sci U S A 117, 28344–28354, doi:10.1073/pnas.2016650117 (2020).

32 Frieman, M. et al. Severe acute respiratory syndrome coronavirus ORF6 antagonizes STAT1 function by sequestering nuclear import factors on the rough endoplasmic reticulum/Golgi membrane. J Virol 81, 9812–9824, doi:10.1128/jvi.01012-07 (2007).

33 Du, Y. et al. 3Cpro of foot-and-mouth disease virus antagonizes the interferon signaling pathway by blocking STAT1/STAT2 nuclear translocation. J Virol 88, 4908–4920, doi:10.1128/jvi.03668-13 (2014).

34 Wang, N. et al. Retrospective Multicenter Cohort Study Shows Early Interferon Therapy Is Associated with Favorable Clinical Responses in COVID-19 Patients. Cell Host Microbe 28, 455–464 e452, doi:10.1016/j.chom.2020.07.005 (2020).

35 Lee, J. S. & Shin, E. C. The type I interferon response in COVID-19: implications for treatment. Nat Rev Immunol 20, 585–586, doi:10.1038/s41577-020-00429-3 (2020).

36 Hadjadj, J. et al. Impaired type I interferon activity and inflammatory responses in severe COVID-19 patients. Science 369, 718–724, doi:10.1126/science.abc6027 (2020).

37 Sa Ribero, M., Jouvenet, N., Dreux, M. & Nisole, S. Interplay between SARS-CoV-2 and the type I interferon response. PLoS Pathog 16, e1008737, doi:10.1371/journal.ppat.1008737 (2020).

38 Blanco-Melo, D. et al. Imbalanced Host Response to SARS-CoV-2 Drives Development of COVID-19. Cell 181, 1036–1045 e1039, doi:10.1016/j.cell.2020.04.026 (2020).

39 Felgenhauer, U. et al. Inhibition of SARS-CoV-2 by type I and type III interferons. J Biol Chem 295, 13958–13964, doi:10.1074/jbc.AC120.013788 (2020).

40 Monk, P. D. et al. Safety and efficacy of inhaled nebulised interferon beta-1a (SNG001) for treatment of SARS-CoV-2 infection: a randomised, double-blind, placebo-controlled, phase 2 trial. Lancet Respir Med, doi:10.1016/s2213-2600(20)30511-7 (2020).

41 Glück, S. et al. Innate immune sensing of cytosolic chromatin fragments through cGAS promotes senescence. Nat Cell Biol 19, 1061–1070, doi:10.1038/ncb3586 (2017).

42 Kelley, W. J., Zemans, R. L. & Goldstein, D. R. Cellular senescence: friend or foe to respiratory viral infections? Eur Respir J 56, doi:10.1183/13993003.02708-2020 (2020).

43 Kim, J. A., Seong, R. K. & Shin, O. S. Enhanced Viral Replication by Cellular Replicative Senescence. Immune Netw 16, 286–295, doi:10.4110/in.2016.16.5.286 (2016).

44 Menachery, V. D. et al. Middle East Respiratory Syndrome Coronavirus Nonstructural Protein 16 Is Necessary for Interferon Resistance and Viral Pathogenesis. mSphere 2, doi:10.1128/mSphere.00346-17 (2017).

45 Zhang, Y. Y., Li, B. R. & Ning, B. T. The Comparative Immunological Characteristics of SARS-CoV, MERS-CoV, and SARS-CoV-2 Coronavirus Infections. Front Immunol 11, 2033, doi:10.3389/fimmu.2020.02033 (2020).

46 Corman, V. M. et al. Detection of 2019 novel coronavirus (2019-nCoV) by real-time RT-PCR. Euro Surveill 25, doi:10.2807/1560-7917.es.2020.25.3.2000045 (2020).

